# A bistable orthogonal prokaryotic differentiation system underlying development of conjugative transfer competence

**DOI:** 10.1101/2021.11.30.470536

**Authors:** Sandra Sulser, Andrea Vucicevic, Veronica Bellini, Roxane Moritz, François Delavat, Vladimir Sentchilo, Nicolas Carraro, Jan Roelof van der Meer

**Affiliations:** Department of Fundamental Microbiology, University of Lausanne, Switzerland

**Keywords:** integrative and conjugative element, Pseudomonas, horizontal gene transfer, regulon, evolution, single cell phenotypic heterogeneity

## Abstract

The mechanisms and impact of horizontal gene transfer processes to distribute gene functions with potential adaptive benefit among prokaryotes have been well documented. In contrast, little is known about the life-style of mobile elements mediating horizontal gene transfer, whereas this is the ultimate determinant for their transfer fitness. Here, we investigate the life-style of an integrative and conjugative element (ICE) within the genus *Pseudomonas* that stands model for a widespread family transmitting genes for xenobiotic compound metabolism and antibiotic resistances. The ICE only transfers from a small fraction of cells in a population, which we uncover here, results from a dedicated transfer competence program imposed by the ICE. Transfer competence is orthogonally maintained in individual cells in which it is activated, making them the centerpiece of ICE conjugation. The components mediating transfer competence are widely conserved, underscoring their selected fitness for efficient transfer of this class of mobile elements.

## Introduction

Prokaryote genomes typically contain a variety of mobile genetic elements (MGEs), such as plasmids^1^, phages, transposons or integrative and conjugative elements (ICEs)^2,3^, which largely contribute to host evolution and community-wide adaptation through genome rearrangements and horizontal gene transfer^4–6^. Although transfer mechanisms per se are well understood, it is insufficiently appreciated that MGEs form their own entities embedded within the host, undergoing selection towards their own fitness optimization^7–11^; for example, by increasing transfer success to new cells^12,13^. MGE decisions potentially oppose the interests of the host cell^14,15^ and can inflict serious damage by cell lysis^16,17^ or cell division inhibition^18–20^. In order to operate independently, MGE regulatory networks need to impinge on the host, but maintain orthogonal components to pursue their own program. Apart from bacteriophage development^21,22^, it is mostly unknown if and how MGEs achieve orthogonality^23^. In order to study fitness selection and orthogonality in horizontal gene transfer, we focus here on a class of mobile DNA elements called ICEs. ICEs have elaborate regulatory systems that control their excision and transfer in response to environmental and host cell cues^24^. They come in various families with distinct and mosaic evolutionary origins. The *clc* integrative and conjugative element in *Pseudomonas* (ICE*clc*) that we use here, stands model for a widespread family occurring in opportunistic bacteria including pathogenic *Pseudomonas aeruginosa*^25,26^. ICE*clc*-type elements have been implicated in transmission of antibiotic resistances^27–29^ and xenobiotic metabolism^25,30^, lending broad significance for understanding the molecular and regulatory basis of their evolutionary success.

ICE*clc* is 103 kb in size and present in two integrated identical copies in the bacterium *P. knackmussii* strain B13^25^, an organism isolated for its capacity to grow on the xenobiotic compound 3-chlorobenzoate (3CBA)^31^. Characteristic for ICE*clc* is its activation in stationary phase cells^32^, leading to excision^13^, temporary replication^12^ and transfer of the ICE^13^ (Fig. 1a). Activated cells are set apart by expression of the promoter of the ICE*clc* integrase (P_int_) and of the *integrase regulatory factor* gene *inrR*^33^. Previously, we showed that transfer of ICE is mediated from a few percent of individual cells in stationary phase that we called transfer competent (tc)^12,13,19^, which is initiated by the activation of a bistable switch^26^. If indeed solely tc cells would be responsible for ICE transfer, it would imply that they (and only they) would have to coherently follow the complete ICE*clc* activation program. This would then suggest the ICE to initiate and orchestrate a specific program in a subset of cells. The nature of the transfer competence program, its temporal and orthogonal coordination in individual cells has remained elusive and is the main focus of this work.

**Figure 1.**
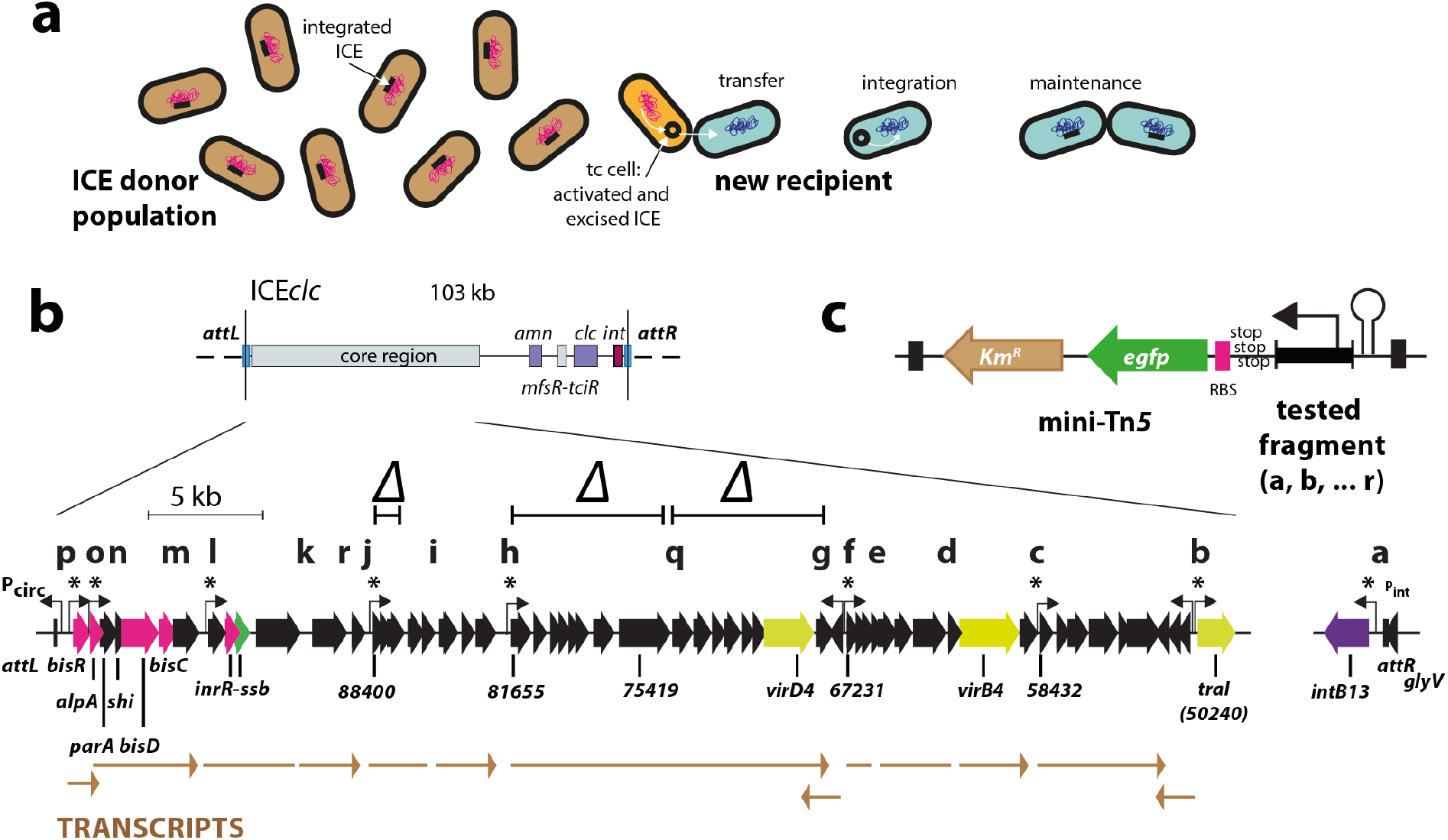
ICE*clc* life-style and transfer competence development. **a** Chromosomally-integrated ICE (black bar, schematically) activates and excises in rare transfer competent cells (tc, black circle; cell highlighted in orange), and transfers by conjugation to a new recipient (here in blue shading), where it inserts site-specifically and maintains through chromosomal replication. **b** Integrated ICE*clc* is delineated between the two attachment (*att*) sites (vertical lines). Layout shows the location of the core region relative to the integrase gene, the variable gene region with the *clc* genes for chlorocatechol and *amn* for 2-aminophenol metabolism, and the key regulator genes *mfsR-tciR*. Gene map below shows individual core genes (black and colored arrows, relevant gene names underneath), previously Northern-mapped transcripts and their orientation (in brown), fragments tested for promoter studies (letters on top) and hooked arrows pointing to identified promoters. Asterisks point to those promoters being expressed in tc cell subpopulations. Regions indicated with a Δ denote deletions for functional studies. **c** General strategy of single copy chromosomally delivered individual or paired promoter-fluorescent gene reporter fusions using mini-transposon delivery.

ICE*clc* transfer competence is most likely encoded by a core set of ~40 genes (Fig. 1b) that is highly conserved among a wide variety of *Proteobacteria*^25,30^. Bistable activation of the ICE in a subpopulation of cells involves three different regulatory loci acting sequentially and initiating a self-regulatory feedback loop^26^ under stationary phase conditions^32^. The role of the feedback loop was suspected to provide a constant supply of a transcription activator complex named BisDC^26^, which we hypothesise, coordinates activation of the different core gene components needed for ICE excision and transfer, all taking place in the same subpopulation of tc cells. In order to study this hypothesis, we selected all potential promoter regions from transcriptional units that were previously described within the conserved ICE*clc* core region^34^, and examined their temporal and subpopulation expression from single-copy chromosomally integrated fusions to fluorescent reporter genes in *Pseudomonas* carrying ICE*clc*, either alone or in combinations. We verified the dependency of identified promoters on key ICE*clc* regulatory elements, using single-cell fluorescence imaging and RNA-seq, and studied functionality of implied gene regions in ICE transfer and their conservation in other ICE of the same family. Our results indicate a multi-operon bistable regulatory network of transfer competence formation that is followed by and restricted to a subset of cells, assuring orthogonality and streamlined ICE*clc* transfer.

## RESULTS

### Identification of subpopulation-expressed promoters within the conserved core region of ICE*clc*

To identify the gene regions possibly implicated in the ICE*clc* transfer competence regulon, we inspected all the putative promoters in the conserved core region of ICE*clc* (Fig. 1b). Putative promoter regions were selected based on a previously conducted ICE*clc* transcript analysis (Fig. 1b, brown arrows)^34^ and further verified by global transcript analysis (RNA-seq, see below). Eighteen putative promoter fragments were amplified by PCR, fused to a promoterless *egfp* gene with its own ribosome binding site (Fig. 1c, Supplementary figure S1), and placed in single copy on the chromosome of *P. knackmussii* or *Pseudomonas putida* containing the ICE. Fluorescent protein expression was examined in stationary phase cultures grown on 3CBA for three clones of each fusion construct, inserted at different random chromosomal positions, in order to control for single-copy positional effects. We looked at both generally increased cellular fluorescence as well as subpopulation-specific increases.

Nine cloned regions yielded clear eGFP expression in stationary phase cells, with a typical small proportion of brighter cells amidst the rest (Fig. 2a). Their bimodal expression becomes more apparent from quantile-quantile analysis, yielding two separate population distributions, the largest of which with low baseline expression and the smaller with distinct higher eGFP expression (white and yellow zones, respectively, Fig. 2b). This contrasts to the eGFP fluorescence distributions observed with the other tested regions, which did not show any deviation from a single expected unimodal distribution (Fig. 2c). The average expression of the identified subpopulations across three independent clones with different single-copy insertion positions was significantly higher for the nine promoter regions than the mean eGFP fluorescence of their main populations (compare magenta to brown bars, Fig. 2d, *n* = 3 replicates, p-values from paired t-tests). For none of the other fragments in any of the three clones tested, subpopulations of cells with higher eGFP expression were apparent (Fig. 2d). Some differences in the mean fluorescence levels of their main cell populations were visible, some being higher (i.e., UR-parA, UR97571, UR89247 and UR73676) than a background control of a strain carrying a deletion in a subpopulation-dependent expressed promoter (i.e., 50240Δ, Fig. 2c and d); others no different from background (e.g., UR89746, UR84835, UR66202 and UR62755). These results suggested that nine regions comprise ICE*clc* promoters with bimodal behaviour (highlighted with asterisks in Fig. 2d), whereas the other nine do not. RNA-seq analysis (see below) suggests some of those to have weak but global promoter activity (e.g., the 67800-region). Of the nine ‘bimodal’ promoters, the previously described P_inR_- and P_int_ promoters^33^ were the strongest, and P_88400_ and P_alpA_ the weakest (Fig. 2b and d). The average size of the stationary phase subpopulation with higher eGFP expression (defined by the qq-plot threshold and visualized by the yellow zone in Fig. 2c) varied between 1.5 and 7.5% (Table 1). The highest subpopulation of cells was detected for the *bisR* promoter, which expressed in 9.1% of cells (Table 1).

**Figure 2.**
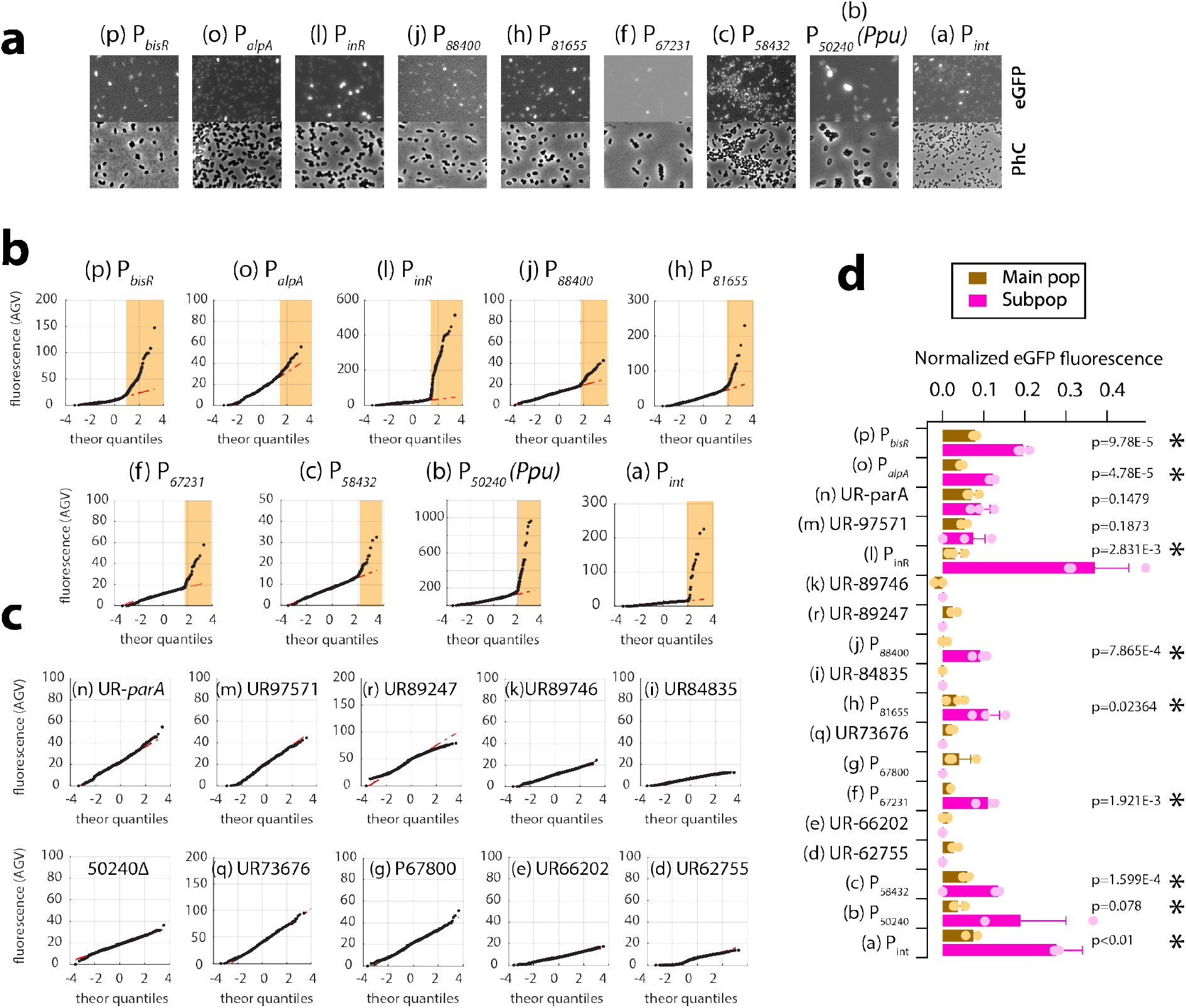
Identification of tc cell subpopulation-specific ICE*clc* promoters. **a** Auto-contrasted and cropped micrographs of *P. knackmussii* B13 strains (exception construct b in *P. putida* ICE*clc*, Ppu) grown on 3CBA imaged in stationary phase for eGFP fluorescence of the indicated construct or in phase-contrast (PhC). White bars indicate 1 μm length. Letters and fragments corresponding to Figure 1 locations. P, putative promoter; UR, upstream region. **b** Quantile-quantile plot representations of stationary phase reporter fluorescence (each dot is value from a single cell). Yellow zones point to tc cell subpopulations. Fluorescence values scaled to zero by subtracting image background. **c** As (**b**) but for the tested constructs without detectable subpopulation expression. **d** Mean normalized fluorescence values (bars) ± one *SD* (whiskers) from 3 replicate strains of *P. knackmussii* B13 or *P. putida* ICE*clc* (construct b) in stationary phase on 3CBA with independent mini-transposon-inserted reporter constructs. Brown bars, value of the main population; magenta bars, values of the identified tc cell subpopulation (if any). Colored dots, individual replicate values. Asterisks point to those constructs showing significantly higher promoter expression in the subpopulation than in the main population (*n* = 3 or 4, p-values from paired one-sided t-test).

**Table 1.** Proportion of stationary phase *P. knackmussii* B13 or *P. putida* ICE*clc* cells expressing eGFP fluorescence from single-copy integrated bistable ICE*clc* promoter reporter fusions

| Promoter region | Wild-type ICE*clc*<sup>a</sup> | Δ*mfsR*<sup>b</sup> |
| --- | --- | --- |
| *bisR* | 9.10 ± 0.87 | 100 |
| *alpA* | 3.10 ± 1.07 | 49.4 ± 2.5 |
| *inrR* | 7.51 ± 1.78 | 77.1 ± 22.0 |
| *orf88400* | 2.30 ± 0.13 | 16.4 ± 2.0 |
| *orf81655* | 1.47 ± 0.47 | 42.2 ± 1.5 |
| *orf67231* | 7.03 ± 2.22 | 40.1 ± 12.3 |
| *orf58432* | 1.59 ± 0.37 | ND<sup>d</sup> |
| *orf50240* | 2.21 ± 0.70<sup>c</sup> | ND |
| *intB13* | 4.7 ± 1.4<sup>e</sup> | 79.0 ± 17.9 |

a) Mean proportion ± one SD of *P. knackmussii* B13 cells expressing eGFP from the corresponding promoter reporter fusion, after 72 h of growth in MM with 5 mM 3CBA, determined by quantile-quantile-plotting (n = 3 independent clones).
b) As a, but in *P. putida* UWC1 ICE*clc*- $\Delta$ *mfsR*.
c) As a, but in *P. putida* UWC1 ICE*clc*.
d) ND, not determined
e) Data reproduced from ref<sup>58</sup>.

Inspection of read coverages from RNA-seq of *P. putida* cells carrying ICE*clc* in exponential growth on 3CBA and subsequent stationary phase (Fig. 3), and in stationary phase of succinate-grown cells (Supplementary figure S2) broadly confirmed specific transcription initiation of the suspected promoter regions. Read coverages in the upstream regions of the open reading frames (ORFs) 67231, 81655, 88400, *alpA* and *inrR* clearly and strongly increased in stationary phase compared to exponential growth on 3CBA (Fig. 3) and in 3CBA versus succinate stationary phase (Supplementary figure S2). The region upstream of ORF58432 showed read-through arising from upstream-located genes (Fig. 3) that may have contributed to its weaker promoter activity in isolation (Fig. 2b). Upstream regions of the ORFs 73676, 84835 and 89247 also showed increases in coverage in 3CBA-grown stationary phase cells (Fig. 3, Supplementary figure S2), but this was not associated with stationary phase expression of single-copy ectopically placed fragments transcriptionally fused to promoterless *egfp* (Fig. 2d). We cannot exclude that the fragments we cloned were too small or otherwise did not encompass the complete promoter region. Alternatively, the observed changes in read coverage in these three regions may have been due to transcript processing as previously suspected^34^. RNA-seq did not support any promoter presence for the regions UR62755, UR66202 and UR97571 (Fig. 3), confirming single cell reporter results (Fig. 2c). Read coverage abundances from RNA-seq did not necessarily correlate to the measured eGFP fluorescence level from cloned fragments in single cells. For example, the read coverage upstream of ORF81655 was the highest of all (Fig. 3); yet the P_81655_-*egfp* fusion was not the highest expressed (Fig. 2c), suggesting additional post-transcriptional control mechanisms or promoter titration effects.

**Figure 3.**
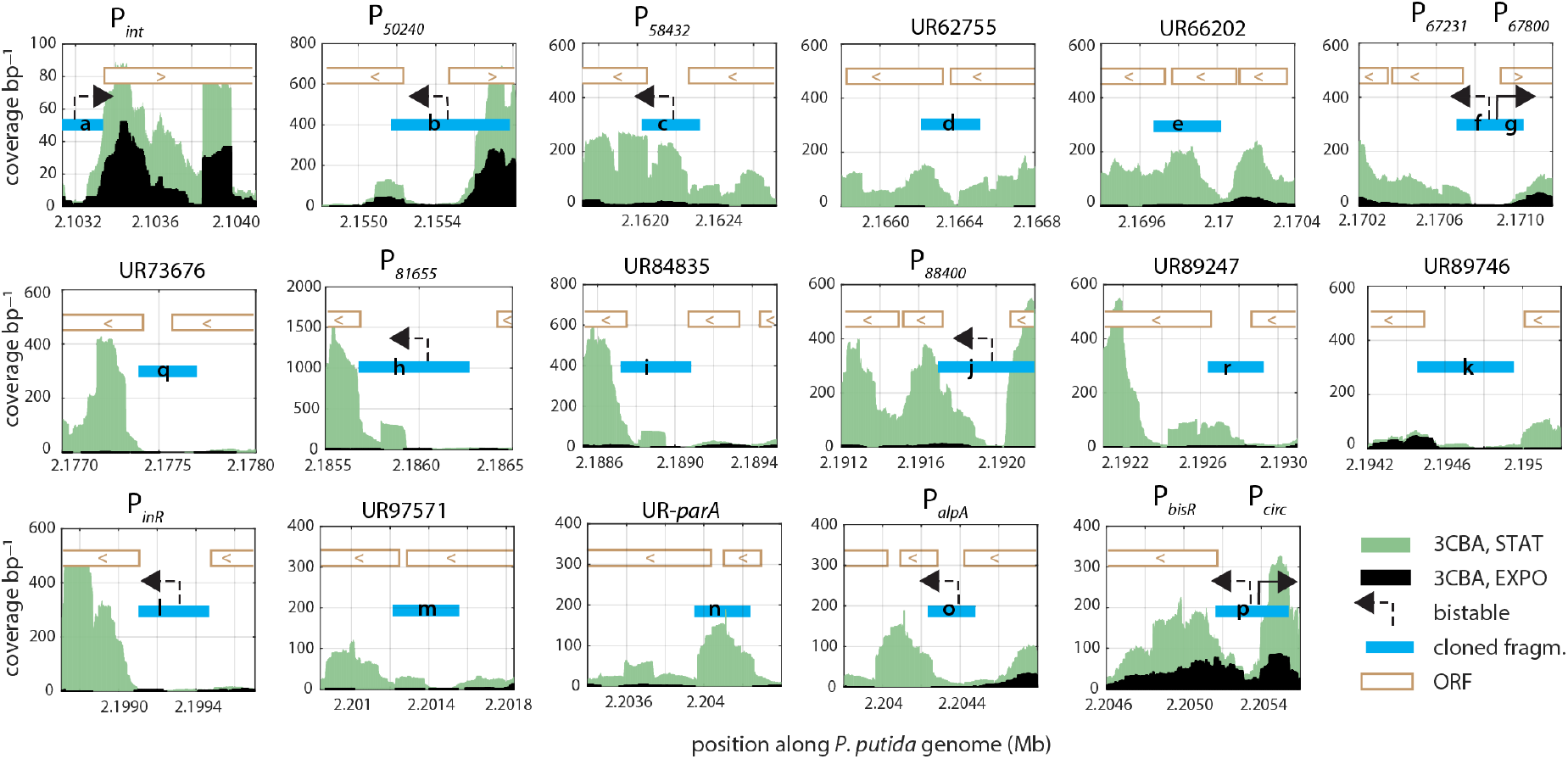
Read coverage of ICE*clc* transcription in *P. putida*. Plots show regions tested for promoter activity with read coverage per basepair position from RNA-seq (for a single representative replicate) at the indicated conditions (3CBA, exponential phase in black; stationary phase in green), plotted for the relevant *P. putida* genome region with the integrated ICE*clc* on the x-axis (in Mbp). Blue lettered bars point to cloned fragments (of Figure 1) tested for promoter activity at single cell level. Dotted black arrows point to subpopulation-dependent tc cell promoters; straight lines when expressed in all cells. Open directional bars (< or >) correspond to relevant coding regions on ICE*clc*. P_circ_, outward facing constitutive promoter^65^.

Read coverages in the upstream regions of *bisR, intB13* and ORF50240 differed much less between 3CBA-stationary phase and exponentially growing or succinate-grown stationary phase cells, suggesting different additional regulatory mechanisms (Fig. 3). However, it should be noted that RNA-seq captures an average from all cells in culture and, therefore, does not exclusively quantify transcripts in subpopulations of transfer competent cells. Coverage plots also suggested specific transcription from the ORFs oriented oppositely to ORF50240 (ORF52324) and from ORF67800, indicative for promoters that would be independent from 3CBA stationary phase conditions (Fig. 3 and Supplementary figure S2).

Collectively, these results indicated that a total of nine upstream regions of the ICE*clc* core genes showed bimodal expression in a subpopulation of stationary phase cells after growth on 3CBA, which suggested they may constitute a coherent regulon.

### Bimodally expressing promoters and P_int_ expression colocalize in the same subpopulations of cells

In order to further determine whether the identified ICE*clc* promoters might belong to a unique concerted regulon operating in the same individual cells, we compared their expression from single copy fluorescent reporter insertions with that of a single-copy P_int_-*mcherry* fusion in the same cell (placed in the same chromosomal position for all comparisons). Stationary phase expression in 3CBA-grown cultures was clearly correlated to that of P_int_ in the case of the P_alpA_, P_inR_, P_88400_, P_81655_, and P_67231_ promoters (Fig. 4a–e, 63.3–91.1% overlap of marker expression in the identified subpopulations, with between 1.7-4.2% of cells being part of the higher-expression subpopulation). Correlation was less clear for the combination of P_int_ with P_58432_-*egfp* (Fig. 4f, 30.4% of marker overlap), which might be due to its inherently lower expression (e.g., Fig. 2b). For unknown reasons, the combination of P_int_ with P_50240_-*egfp* only expressed both markers in *P. putida* ICE*clc* but not in *P. knackmussii* B13, with slightly lower subpopulations of cells than before (Fig. 4g, 0.8-2.0% and 54.8% of marker overlap). Interestingly, the combination of P_int_ to P_bisR_ only showed 13.8% of overlap in cells with both subpopulation-expressed markers, and a larger subpopulation of P_bisR_-eGFP-expressing cells than for the other constructs (Fig. 4h, 6.1%, as noted before for the individual promoter fusions, Table 1). These results thus suggested that, apart from the *bisR*-promoter, the other bimodal ICE-promoters express in the same individual cells.

**Figure 4.**
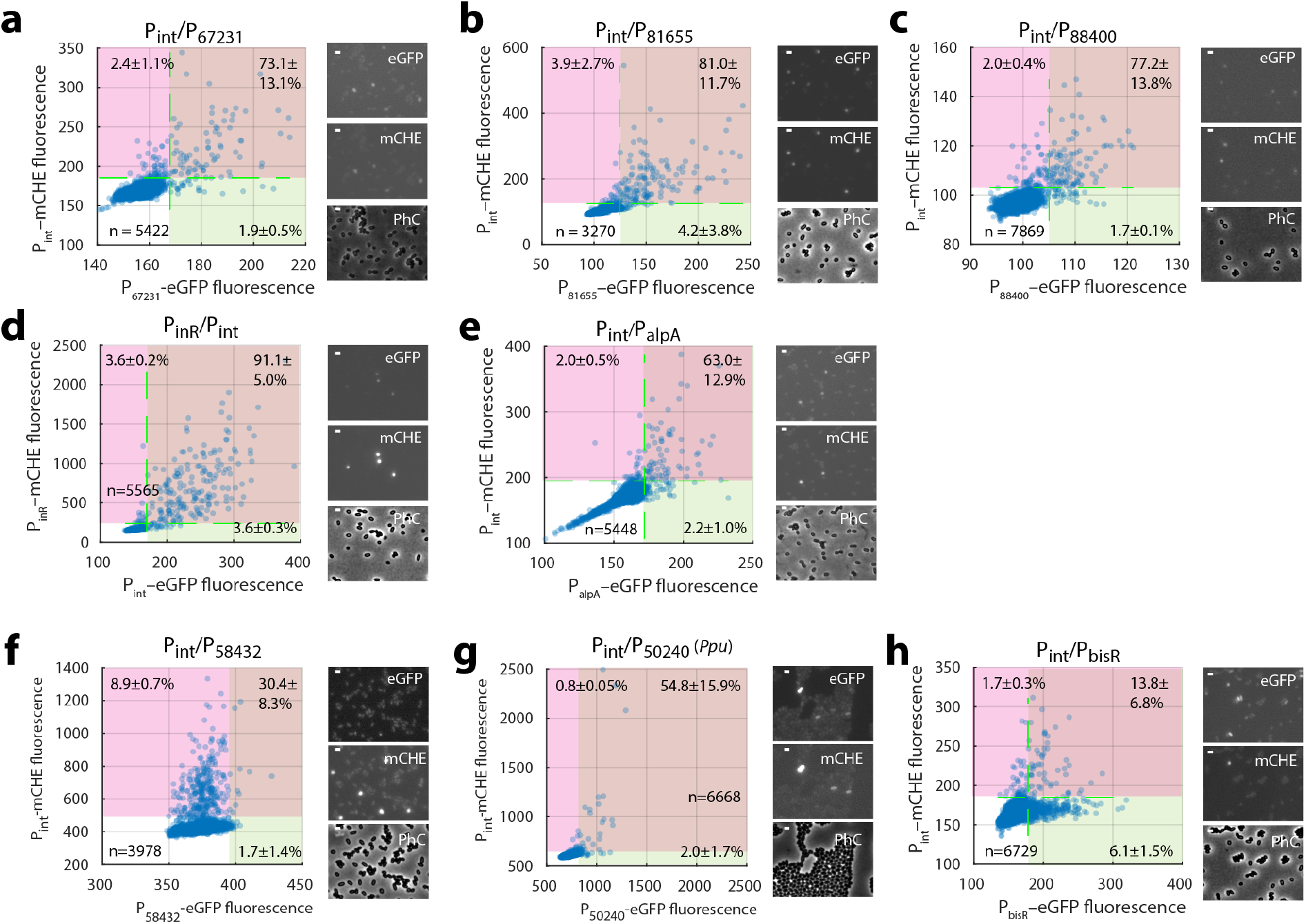
Colocalization of expression of paired bimodal promoters from the ICE*clc* transfer competence regulon (**a-h**). Dots show stationary phase fluorescence values (72-h-cultures on 3CBA) of individual *P. knackmussii* B13 or *P. putida* ICE*clc* (Ppu in **g**) cells carrying the indicated (single copy integrated) double reporter construct. Plots combined from three biological replicates (*n* is total amount of analyzed cells) with micrographs on the right showing example images in GFP, mCherry and phase-contrast (auto-contrasted for display). Magenta zones indicate the subpopulation expression in the mCherry-channel (defined from quantile-quantile plots as in Figure 2b, percentages showing the mean subpopulation size ± one *SD* from *n* = 3 biological replicates). Green zones represent the same but for the GFP-channel. Percentages on the upper right indicate the mean proportion ± one *sd* of cells in the overlap (brown) of green and magenta channels. Bars within micrographs indicate 1 μm.

To understand how different temporal expression would influence observed marker correlations in cell populations, we next quantified dynamic reporter fusion expression in growing and stationary phase cells for a subset of the double-labeled strains (Fig. 5). We excluded the P_58432_- and P_50240_-fragments from this analysis, because of their poorer correlations with P_int_-expression observed above. The example of *P. knackmussii* B13 tagged with both P_81655_-*egfp*/P_int_-*mcherry* shows clearly how only a subset of cells fires both promoters after the onset of stationary phase (>8 h, Fig. 5a). Similar behaviour was shown by strains labelled with P_inR_-*egfp*, P_88400_-*egfp* or P_alpA_-*egfp*, and P_int_-*mcherry* (Supplementary figure S3). In contrast, *P. knackmussii* B13 tagged with both P_bisR_-*egfp*/P_int_-*mcherry* lacked coordination of expressing eGFP and mCherry (Fig. 5b), with a subset of cells expressing only eGFP from P_bisR_ but no mCherry (as suggested by the scatter plot in Fig. 4h; blue lines in Fig. 5b). The absence of co-expression of both markers in most cells was therefore not a consequence of different timing but rather a specific proportion of cells expressing only the *bisR*-promoter. Considering cell-cell variations, the strength of promoter expression was reasonably correlated for both markers, taken as the slope of fluorescence increase after the onset of marker activation in individual cells (r^2^ in between 0.43 – 0.68, Fig. 5c). This could imply a coordinated expression in individual cells. Notable exception to this was again P_bisR_-*egfp*/P_int_-*mcherry*, which showed poor correlation between the expression of both markers (r^2^ = 0.1603, Fig. 5c). Finally, also the timing of expression in individual cells was well correlated between P_int_ and the 81655-, 88400-, *alpA*- or *inrR*-promoters (Fig. 5d), whereas this was less obvious for the combination of the *bisR*- and *intB13*-promoters (Fig. 5d). The majority of P_int_-expression activity occurred between 20–30 h, corresponding to 10–20 h after the onset of stationary phase (Fig. 5e, magenta histograms), which was similar to that of the 81655-, 88400-, *alpA-* or *inrR*-promoters (Fig. 5e, green histograms). In contrast, the majority of cells started expressing the *bisR* promoter construct a few hours earlier (~15 h, or 5 h into stationary phase; Fig. 5e, P_bisR_). In summary, these results thus indicated that the ICE core promoters express mostly in the same cells and at the same moment, which is evidence for them being part of the same transfer competence regulon. Activation of *bisR* is only partly correlated and occurs earlier, probably because, as we will further discuss below, it is one of the key ‘upstream’ regulators for activation of the transfer competence pathway of ICE*clc*^26^. The fact that around half of the cells expressing *bisR* do not subsequently activate the ‘downstream’ P_int_-promoter suggests that it is necessary but not sufficient to continue with full transfer competence.

**Figure 5.**
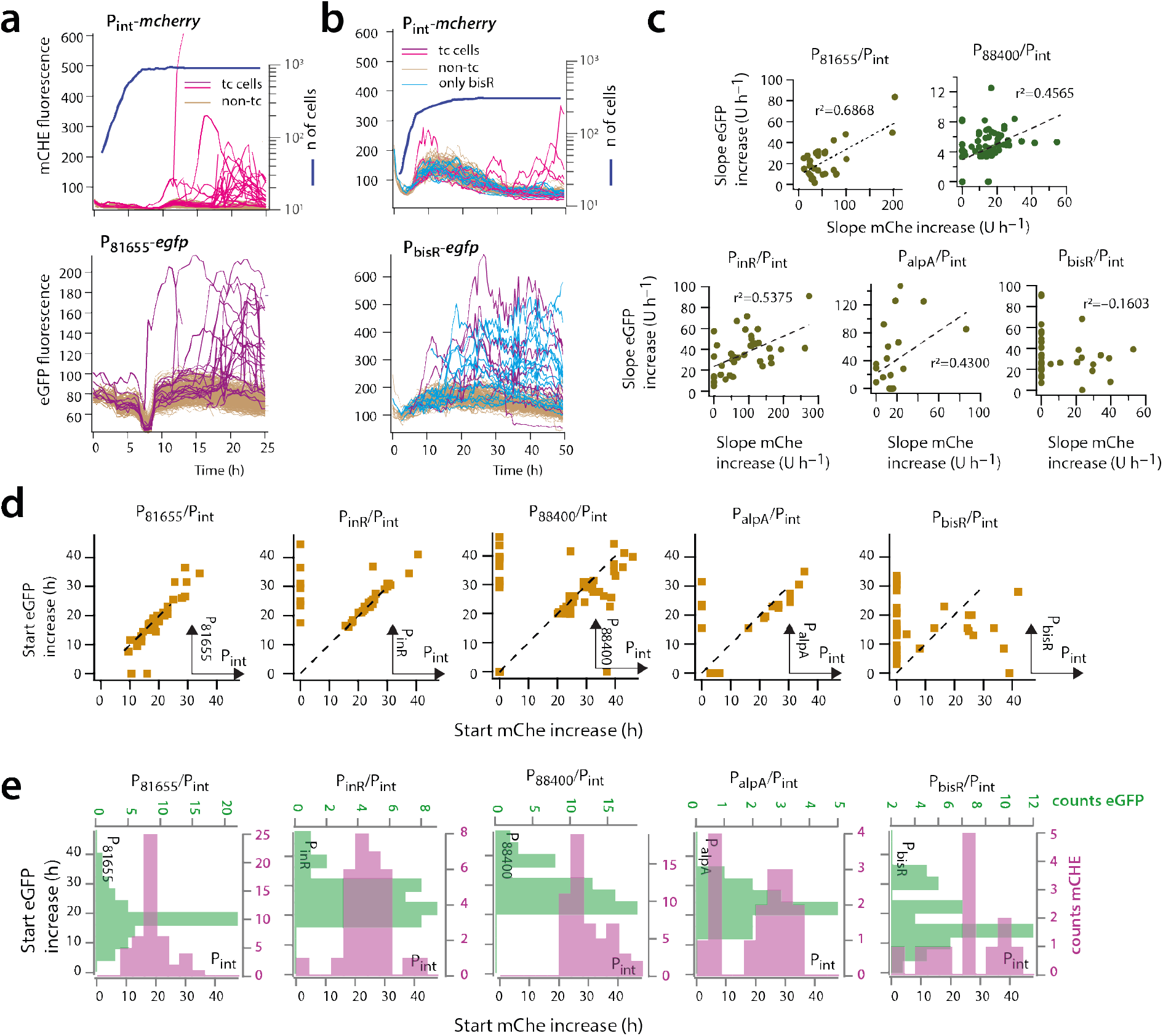
Temporal expression of paired promoters from the ICE*clc* transfer competence regulon. **a** Time-lapse fluorescence of individual cells of surface-grown *P. knackmussii* B13 with indicated single-copy inserted promoter-fluorescent reporter constructs; each line corresponding to an individual cell traced over time. Line traces are colored according to attribution of an individual cell as transfer competent (tc, magenta or purple) or not (brown), based on quantile-quantile plotting at time point 20 h in both channels (plus retracing its previous history). Thick blue line indicates cell growth (scale on the right) for the analyzed image areas. **b** as (**a**), but showing the example of partly uncoordinated expression of the *bisR* promoter firing in twice the amount of cells than the *int* promoter. **c** Correlation of fluorescent reporter expression from paired single-copy inserted constructs of the indicated promoters in *P. knackmussii* cells identified as being tc (i.e., cells with magenta and blue lines in panels **a** and **b**). *int* Promoter in all cases coupled to mCherry. Dots indicate measured slopes of fluorescent signal increase over time (intensity units – U h^−1^), with dotted lines showing linear regression and regression values. **d** Correlations of the onset of marker fluorescence expression in identified tc cells (here as squares) displayed in (**c**), relative to the incubation start (start of stationary phase between 8–10 h, panels **a** and **b**). Dotted lines represent same timing. Dots with ‘zero’ timing have no distinguishable slopes in that channel. **e** Distribution of expression starts for both markers among individual cells of the *P. knackmussii* reporter pairs of (**c**) and (**d**), displayed as histograms for eGFP (green shading) and mCherry (magenta, in all cases from P_int_). Because of slight image drift, the time-lapse data before t=20 h from P_int_/P_88400_ could not be linked.

### Dependency of core promoter expression on known ICE*clc* regulatory factors

In order to confirm whether the identified bistable ICE*clc* promoters would be part of a coherent transfer competence regulon, we examined the dependencies of their expression by RNA-seq and single copy promoter fluorescent reporter fusions in the presence or absence of previously identified key regulatory elements for activation of ICE*clc mfsR*^35^, *inrR*^33^, *bisR* and *bisDC*^26^.

All identified bistable promoters strongly increased their expression in stationary phase *P. putida* ICE*clc* background with a deletion in *mfsR* (Fig. 6a, Δ*mfsR* STAT). MfsR is the major global transcription regulator of ICE*clc* that represses its own transcription and that of *tciR* within the same operon^35^, plus that of a set of genes on ICE*clc* coding for an efflux pump^36^. As previously observed^35^, *mfsR* deletion resulted in increased expression of *tciR* (Fig. 6a, b), and of a set of multidrug efflux pump genes (*mfsABC*, Fig. 6a)^36^. Deregulated *tciR* also led to higher *bisR* expression (Fig. 6a, EXPO), which is linked to ICE*clc* activation^26^. Indeed, global transcript abundances of the ICE*clc* core genes were 2–32 fold higher in stationary phase 3CBA-grown cells of *P. putida* ICE*clc-ΔmfsR* than wild-type ICE*clc* (Fig. 6b). Expression of ORF52324 and ORF67800 was unaffected, confirming they are not part of the bistability regulon (Fig. 6a, b). The increased globally observed expression of the ICE core genes in the Δ*mfsR* strain was primarily due to a sharp increase of the subpopulation of cells activating the ICE compared to wild-type ICE*clc* (Table 1). When additionally to *mfsR* also the *bisR* gene was deleted, stationary phase expression of the ICE*clc* core genes was strongly reduced, even lower than in wild-type (Fig. 6a, b). The reason for this is that in absence of BisR no further activation can proceed^26^. Expression of the *clc* genes in exponential phase remained unaffected, as expected for being outside the transfer competence regulon. That of the MfsR-controlled *tciR* and efflux pump genes^36^ remained constitutive in absence of *bisR* (Fig. 6a), indicating they are not influenced by BisR. Transcription from ORF52324 and ORF67800 was also unaffected by the *bisR* deletion, confirming that they are not part of the transfer competence regulon. Taken together, this indicated that the bistable ICE core promoters are dependent on the early regulators MfsR/TciR, yet only activated through BisR as an intermediate step.

**Figure 6.**
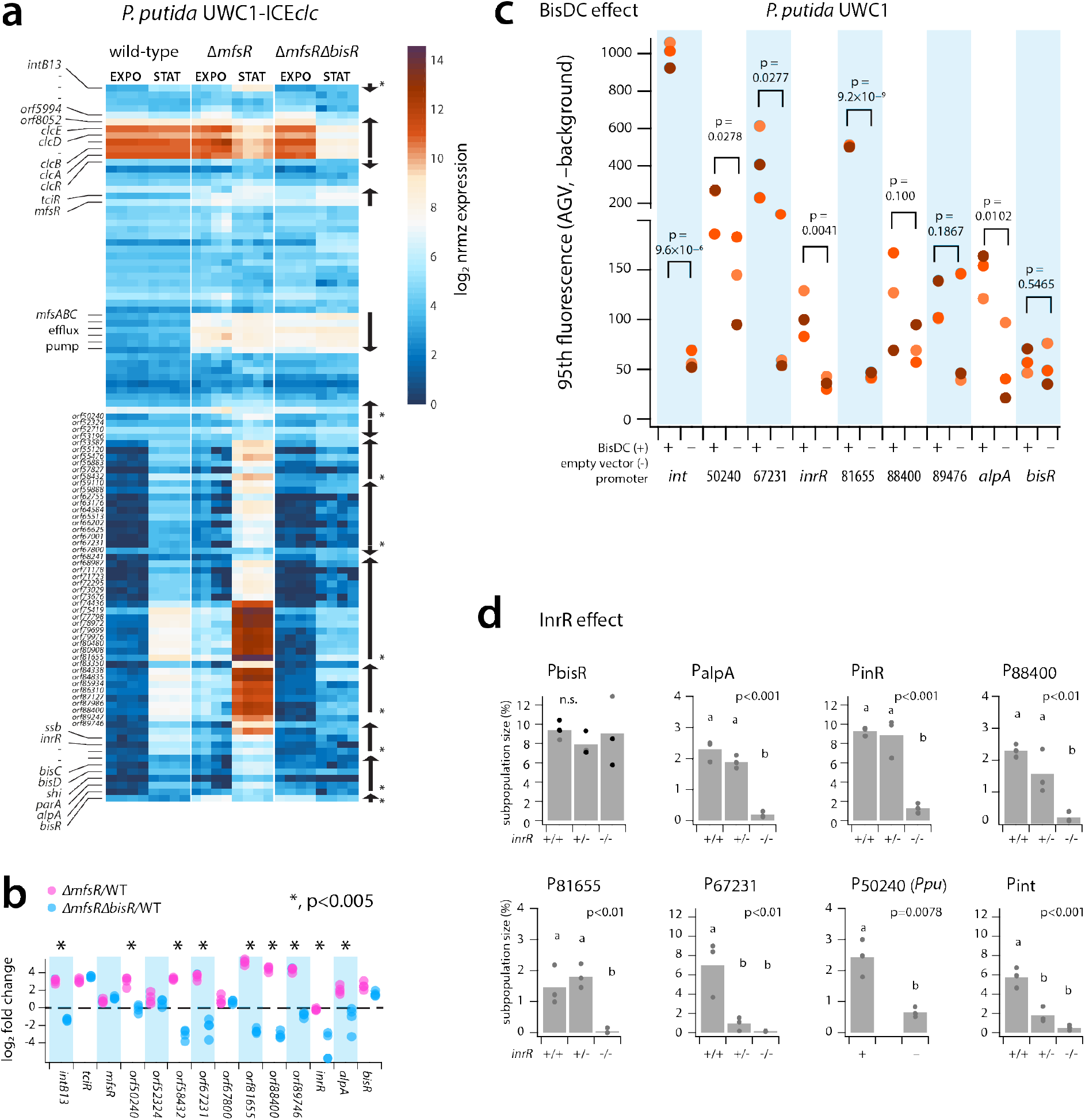
Dependency of ICE*clc* promoters on known ICE global regulators initiating transfer competence. **a** Log_2_-scaled normalized and per-gene attributed read counts from RNA-seq of *P. putida* carrying wild-type ICE*clc*, *ΔmfsR* or *ΔmfsRΔbisR* deletions, for the ICE*clc* gene region (*intB13* gene on top; *bisR* at bottom). Each colored rectangle corresponds to a single gene (relevant names on the left) and replicate (four replicates per condition and strain; growth on 3CBA, sampled in exponential phase and late stationary phase). Organisation of transcriptional units depicted with arrows on the right (asterisks corresponding to those being part of the transfer competence regulon). **b** Calculated log_2_-fold changes between mutant and wild-type ICE*clc* for the tested promoter regions and controls. Dots corresponding to individual replicate values. Dotted line represents ratio of 1. Asterisks denote statistically significant differences in *mattest* with bootstrapping (n = 1000). **c** Effect of induction of plasmid-located *bisDC* (+) on reporter fluorescence of *P. putida* without ICE*clc* with indicated single copy inserted promoter-fusions. Cells sampled in stationary phase after growth on succinate. Comparisons are the same *P. putida* reporter strains but with empty plasmid (–). Plotted is the 95^th^ percentile scaled fluorescence (minus image background) of the population of sampled cells (*n* = 10 technical replicates, each dot from a single biological replicate with independent reporter gene insertion position), to account for the strongly tailed fluorescence distributions. p-values calculated from paired one-sided t-test of strains in presence of BisDC versus the vector-only control (*n* = 3, replicates per construct). **d** Effect of *inrR* (two, one or no copy) on ICE*clc* in *P. knackmussii* B13 or *P. putida* ICE*clc* (P_50240_-construct) on expression of the indicated reporter constructs (in all cases fused to *egfp*, three clones with different single copy insertion positions). Bars show the mean ± one *SD* of the estimated subpopulation sizes (from quantile-quantile plotting) in cultures growing on 3CBA sampled after 24, 48, 72 or 96 h (24 h corresponding to onset of stationary phase). Letters correspond to significance values in one-factorial ANOVA across all samples followed by post-hoc Tukey test.

Next we tested whether the suspected ICE*clc* promoters could be directly activated in absence of the ICE in *P. putida* by overexpression of BisDC, which is the previously identified key regulator for controlling and maintaining bistable expression^26^. IPTG induction of plasmid-localised *bisDC* expression in *P. putida* without ICE*clc* but equipped with single copy inserted promoter-fused fluorescent gene reporters resulted for all tested constructs in increased reporter fluorescence compared to an empty plasmid control (Fig. 6c), except for the negative control (UR89476) and for the *bisR* promoter. The magnitudes of expression, however, were very different among the tested promoters and several resulted in highly skewed or subpopulation-confined expression (hence testing the 95^th^ percentile levels in Fig. 6c; Supplementary figure 4). These results thus demonstrated that these promoters are ‘downstream’ in the regulatory cascade of *bisR* and the *bisDC* feedback loop. Finally, we tested whether their expression was dependent on InrR, a previously reported factor contributing to optimal expression of P_int_^33^. For this, we introduced the single copy promoter fluorescent reporter gene fusions in a *P. knackmussii* wild-type background, and with one (*inrR*^+/-^) or both copies of *inrR* deleted (*inrR^-/-^*), and measured subpopulation-dependent fluorescence expression in stationary phase cells of 3CBA-grown cultures (Fig. 6d). For unknown reasons, the ORF50240-promoter construct did not express in *P. knackmussii* B13 and was again tested in *P. putida* with one ICE*clc* copy. Apart from P_bisR_, all other promoters were dependent on InrR, with subpopulation sizes strongly diminished in absence of one or both *inrR* gene copies (Fig. 6d). Collectively, these results thus indicated that the ICE*clc* transfer competence regulon encompasses a number of ‘late’ expressed elements (promoters upstream of ORF88400, ORF81655, ORF67231, ORF58432, ORF50240, and *intB13*). Their expression is dependent on factors produced in the early stages (e.g., TciR, BisR) and from the feedback loop (BisDC, InrR).

In order to get an idea on the possible functional importance of ICE genes expressed from the late promoters, we produced a number of seamless deletions on the ICE*clc* and tested their effect on activation and transfer rates. Deletion of the region from ORF81655-75419 or from ORF88400-84388 had no measurable effect on transfer from *P. putida* to a gentamicin-resistant isogenic strain, whereas that of ORF74436-68241 abolished transfer completely (Fig. 7). All constructs expressed similar proportions of tc cells in stationary phase (Fig. 7), indicating that absence of transfer was not due to abolished induction of transfer competence.

**Figure 7.**
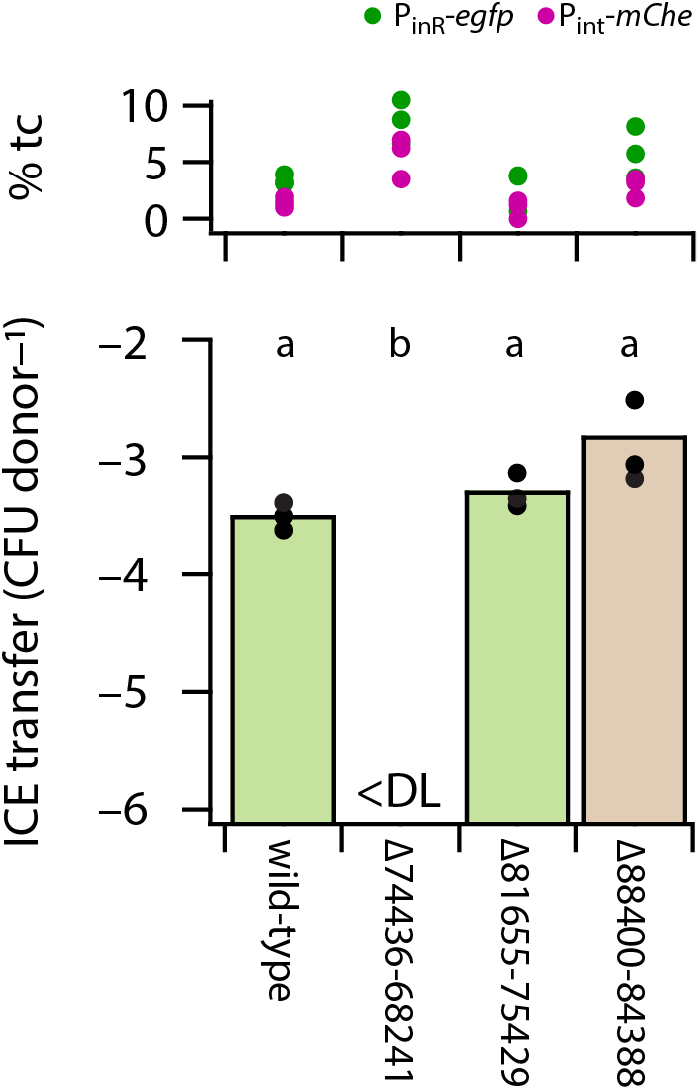
Effect of ICE*clc* gene region deletions on transfer frequency (bottom) and subpopulation size of P_int_-mCherry/P_inR_-eGFP expressing cells (top, as percentage from qq-plots). Bars show the calculated mean ICE transfer frequency (as colony forming units of transconjugants per CFU of the donor) for the indicated ICE in *P. putida* (deletions as in Figure 1a), with dots representing individual replicate values. Letters correspond to significance values (p < 0.005) in one-factorial ANOVA followed by post-hoc Tukey test. <DL, below detection limit.

## DISCUSSION

ICE*clc* had been hypothesized to impose a bistable differentiation program on its host cell that elicits competence for its own transfer^24^. Previous studies visualizing single cell ICE*clc* activation from fused fluorescent protein genes to the integrase promoter P_int_ and the promoter of the integrase regulator gene *inrR* had inferred that this transfer competence would arise in a 3-5% subpopulation of cells carrying the ICE^19,33,37^, under stationary phase conditions^32^ and most pronounced after growth on 3CBA^33,37^. Since ICE conjugative transfer is assumed to involve a variety of distinct steps (e.g., excision, unwinding to single-stranded DNA and replication, presentation to the conjugation complex^24,38^), we hypothesized that transfer competence development would encompass hierarchically and temporally controlled activation of subsets of ICE-genes in the same individual cells that commit to the complete process. We uncovered here that most of the genes within the conserved core region of ICE*clc* belong to a program that we can now name ‘transfer competence regulon’ (TCR). The regulon is organized in six ‘downstream’ transcriptional units that are coordinately transcribed within the same subpopulation of cells (Fig. 1 and 8). All downstream transcriptional units are dependent on the transcription activator complex BisDC, and on the previously discovered factor InrR^33^, which are produced in the feedback loop that is initiated by action of BisR and its ‘upstream’ cascade of MfsR and TciR (Fig. 8)^26^. In addition, the core region has two transcriptional units that are oppositely oriented and do not seem to be part of the TCR (i.e., ORF67800 and ORF52324-53196).

**Figure 8.**
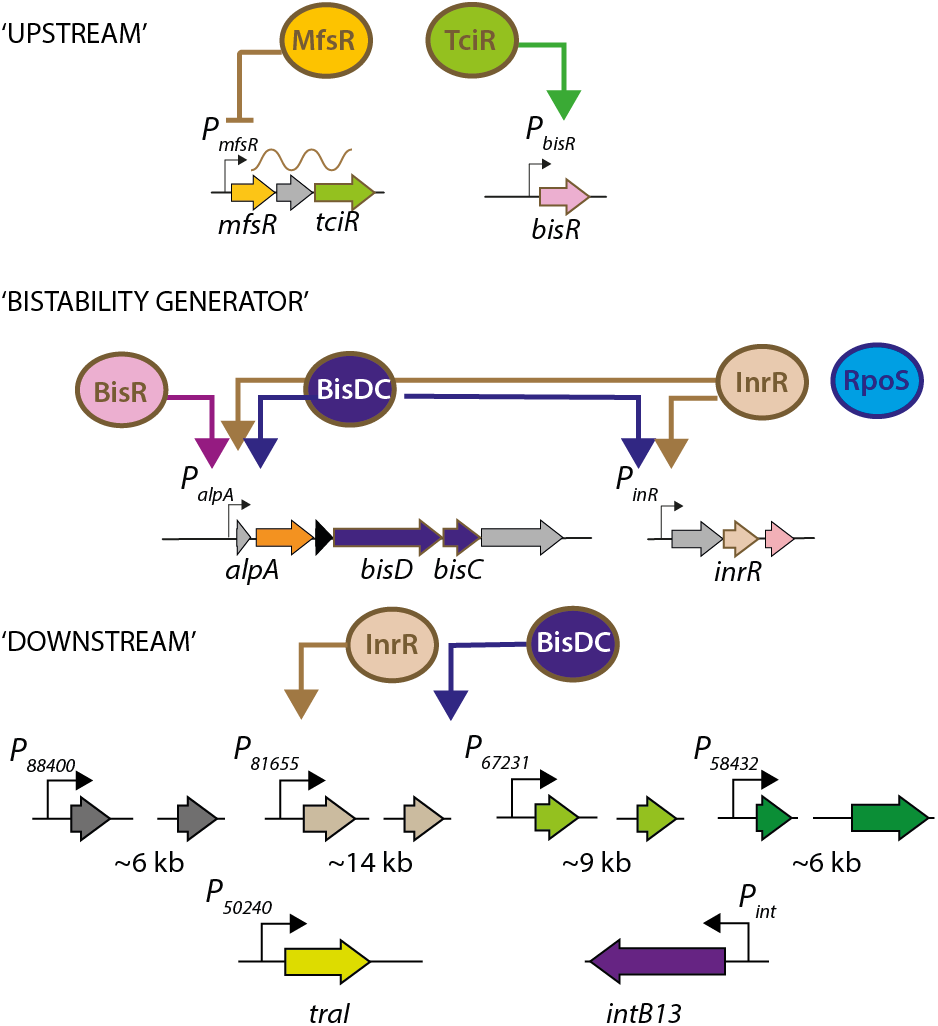
The ICE*clc* transfer competence regulon. Hierarchically ‘upstream’ factors MfsR, TciR and BisR, acting sequentially. The ‘bistability generator’ configuration initiated by BisR, but maintained by BisDC and InrR, under further global control by RpoS^32^. Finally, the ‘downstream’ or ‘late’ targets under orthogonal control of BisDC and InrR, activating in parallel the operons needed for ICE transfer itself.

Our conclusion is based on several lines of evidence, notably RNA-seq and fluorescent reporter expression in single cells controlled from different individual or pairs of ICE core promoter regions, in ICE*clc* wild-type and mutant backgrounds. RNA-seq helped to identify global regulatory effects and transcript differences, and the read coverage abundances supported attribution of potential promoter regions. Fluorescent reporter expression was essential to quantify and restrict the subpopulation of stationary phase tc cells, which is a hallmark of the ICE activation program^33^. Although we may have missed minor promoter details by cloning, we are fairly confident to have uncovered the major transcriptional units of the TCR and their attribution in the expression hierarchy (Fig. 8). Furthermore, although taken at face-value the expression of each of the individual TCR promoters is bimodal within the population of stationary phase cells, pair-wise and temporal expression patterns indicate them to transcribe in the same individual cells, thereby forming a single bistable pathway. We thus firmly show that the TCR, once initiated, imposes a bistable developmental pathway on a subset of cells only. This pathway leads to cells being able to excise, process and transfer the ICE, and to them arresting cell division upon renewed growth, as was shown elsewhere^18,19^. This also fulfills the second requirement of a bistable pathway, namely, the state of no-return^39–42^.

Among the late genes of the TCR, gene deletion and mating experiments suggested that at least the second half of the multicistronic unit downstream of the P_81655_ promoter is essential for ICE*clc* transfer, whereas genes included in the regions under control of P_67213_ and P_58432_ may code for distant type IV secretion system components^34^; and, thus, would be essential for ICE transfer as well. The role of the relaxase gene (*orf50240*) for ICE*clc* had been shown previously^43^ and we confirm here that its expression belongs to the TCR. By contrast, the role of the genes under control of the P_88400_ promoter for ICE transfer is not clear, since their deletion did not result in measurable reduction of transfer frequency from a *P. putida* ICE donor to an isogenic *P. putida* recipient. The genes in this region code for conserved hypothetical proteins leaving little room for speculation as to their potential function in ICE*clc* transfer. However, many of them are conserved among ICEs of the same family in different hosts both by individual sequence as well as in gene synteny (Supplementary figure 5), suggesting functional importance for ICE maintenance, regulation and/or transfer.

One of the intriguing questions in the ICE*clc* TCR pathway is to understand how its orthogonality can be achieved. In other words: how can be guaranteed that individual cells, which initiate the pathway, will also continue along its path and successfully transfer the ICE? And, secondly: how can it be avoided that parts of the TCR pathway are expressed in non-tc cells that might impact their survival (given that TCR eventually comes with a cost of cell arrest^18,19^)? Bacteriophages have solved the problem of orthogonal expression of components, for example, by coding for their specific phage DNA and RNA polymerases^21,22^. As the ICE does not appear to encode such RNA polymerase it needs to accomplish this task differently, potentially by maintaining control of the promoters through two factors (BisDC and InrR), which are both exclusively expressed in tc cells. Time-lapse single cell expression data from individual and pairs of TCR promoters indicated that activation is quite noisy, and is very sensitive to placement of additional promoter copies (i.e., apart from the ICE itself; for example Fig. 5a and b), which is suggestive for promoters regulated by low copy number transcription factors in the cell^44,45^. On the basis of stochastic models, we had previously suggested that the feedback loop (Fig. 8) might function to maintain low but steady levels of BisDC in cells that have initiated the TCR^26^, such that its downstream promoters can be activated. Since this is an autoregulatory loop, changes in “free” BisDC levels as a result of promoter binding, would be compensated for by increased production. On the basis of the results shown here (Fig. 6), we have to conclude that the proposed feedback loop by BisDC may have an additional component that includes InrR, whose biochemical role is so far not understood. Although overexpression of BisDC is sufficient to activate TCR promoters in absence of the ICE, our current working hypothesis is that InrR under wild-type conditions acts in conjunction with BisDC to provide TCR-promoter specific recognition and recruitment of the host RNA polymerase. This would give the fidelity and the orthogonality to the system to follow the transfer competence in the same cells where it initiated.

Another curious discovery here was that expression of the *bisR* promoter is already bimodal, but is visible in twice the proportion of cells that continue the TCR and express the downstream promoters. The same stochastic modeling of ICE*clc* regulation had also suggested that the input levels of BisR at the point of initiating the feedback loop (at the *alpA* promoter) are determinant for its output in terms of proportions of cells with active TCR^26^. Furthermore, a synthetic inducible *bisR* construct produced scalable subpopulation sizes of activated cells^26^. What is then the mechanism that, as we discovered here, subdues BisR activation at *alpA* under wild-type conditions, or rather, seems to lead to abortion of TCR in half of the cells? Without knowing further biochemical details on protein stability and binding constants this is hard to deduce, but possibly also here, the crux lays in the action of InrR as auxilliary protein. We assume, therefore, that it is not only BisDC, but the BisDC/InrR combination that controls the equilibrium of the 2–5% of wild-type cells that develop transfer competence, whereas otherwise the proportion of transfer competent cells would be solely controlled by those expressing BisR^26^.

In summary, we uncovered the extent and the hierarchy of the regulon that encompasses transfer competence development of ICE*clc*, showing how the TCR restricts to this subpopulation of active cells that are the centerpiece of efficient ICE transfer. ICE*clc* has evolved to a remarkably efficient transfer machine, operating within “the window of opportunity” that it creates in a few individual cells to not disturb its host population (too much) and still transfer highly efficiently^24^. Understanding this process and its adaptation is crucial, given the broad occurrence of ICE in prokaryotic genomes^46^, and the particular wide distribution of the ICE*clc* family of elements^25,30^, also among important opportunistic pathogens^47,48^ with ICE-carried antibiotic resistance genes^27–29^. A further central question to solve is the influence of environmental or physiological cues (such as 3CBA metabolism in case of ICE*clc*) on the proportion of appearing tc cells. Such cues or changes in environmental conditions may unwillingly influence gene transfer rates within microbial communities^49–51^, and this may lead to enhanced adaptation of pathogenic isolates to antibiotic resistances carried by the ICE.

## MATERIALS AND METHODS

### Bacterial strains and plasmids

*Escherichia coli* strain DH5α (Gibco Life Technologies, Gaithersburg, Md.) was routinely used for plasmid propagation and cloning experiments. *E. coli* DH5α-λpir was used for the propagation of pBAM plasmids used for the delivery of mini-Tn*5* transposons^52^. The original host harbouring ICE*clc* is *P. knackmussii* B13^25,53^. *P. putida* UWC1 was used as a further host for (a single copy of) ICE*clc*^43^. Bacterial strains, plasmids and primers used in this study are listed in Supplementary tables S1 and S2.

### Media and growth conditions

*E. coli* strains were cultivated in Luria Bertani (LB) medium with incubation overnight (O/N) at 37°C. *P. knackmussii* B13 and *P. putida* were grown at 30°C in minimal medium (MM) based on the type 21C medium^54^ with 5 mM 3CBA or 10 mM succinate as sole carbon and energy source. When required, the following antibiotics were added to the media at the following concentrations: ampicillin 100 μg ml^-1^, kanamycin 50 μg ml^-1^ (*E. coli*) or 25 μg ml^-1^ (*P. knackmussii* and *P. putida*), gentamicin (25 μg ml^-1^), tetracycline 20 μg ml^-1^ (*E. coli*) or 100 μg ml^-1^ (*P. putida*) and chloramphenicol 5 μg ml^-1^. Transcription from the P_tac_ promoter was induced by addition of 0.05 mM isopropyl β-D-1 thiogalactopyranoside (IPTG).

### DNA manipulations

Isolation of chromosomal and plasmid DNA, PCR, restriction enzyme digestion, ligation and electroporation were performed as described by standard procedures^55^, and as previously described^26^. Electrotransformation of *P. knackmussii* and *P. putida* was performed using the procedures described by Miyazaki et al.^43^. Seamless chromosomal deletions on ICE*clc* were produced using I-SceI-induced chromosomal breakage and double recombination, as described^26,56^.

### Reporter gene fusions

Appropriate DNA fragments containing the putative ICE*clc* promoter regions (^34^; Supplementary table 2) were amplified by PCR and cloned in front of a promoterless eGFP gene on the mini-Tn*5* delivery vector pBAM^52^. The resulting pBAM-eGFP promoter reporter fusions were then introduced in single copy onto the chromosome of *P. knackmussii* B13 or *P. putida* UWC1 (ICE*clc*) using electrotransformation. Transformants were selected on Km selective medium and verified by PCR for appropriate integration. For each reporter fusion, at least three independent clones were selected and purified, with which microscopy analysis of eGFP expression was carried out.

For colocalization studies, a mini-Tn*5* containing either of the different promoter-eGFP constructs was inserted in *P. knackmussii* B13 or *P. putida* UWC1 (ICE*clc*) already containing a P_int_-*mcherry* reporter fusion. The resulting transformants were selected on Km and Tc selective medium and verified by specific PCR; and three independent clones were stored.

### Epifluorescence microscopy

For the detection of eGFP and mCherry expression in single cells, *P. knackmussii* B13 and *P. putida* UWC1 strains were cultured for 16 h at 30°C in LB medium. Aliquots of 100 μl of this culture were then diluted in 20 ml MM plus 5 mM 3CBA and corresponding antibiotics, and incubated at 30°C. After 24 h, 48 h, 72 h and 96 h (cultures typically reach the onset of stationary phase between 24 −48 h), a culture aliquot of 400 μl was drawn. The cells in the sample were harvested by centrifugation at 7000 × *g* for 2 min, after which the cell pellet was carefully resuspended in 50 μl of fresh MM without carbon source added. An aliquot of 4 μl suspension was then spread onto a regular microscopy slide, precoated with 0.7 ml of a 1% agarose in MM solution. Slides were covered with a 50 mm x 15 mm cover slip, moved to the dark room, after which cells were imaged (typically within 15–30 min after application; aiming to have between 5–10 images with *n* = 1000 cells in total per clone and time point). Images were taken under phase-contrast (10 ms), eGFP fluorescence (500 ms), and mCherry (500 ms) using a Zeiss Axioplan II imaging microscope with a 100× Plan Apochromat oil objective lens (Carl Zeiss, Jena Germany) and equipped with a SOLA SE light engine (Lumencor, USA). A SPOT Xplorer slow-can cooled charge coupled device CCD camera system (1.4 Mpixel; Diagnostic Instruments, Sterling Heights, Mich.) fixed on the microscope was used to capture the images. Cells on images were automatically segmented using SuperSegger^57^ as previously described^12^, calculating average per cell fluorescence intensities in the eGFP and/or mCherry channels. Subpopulations of tc cells were inferred and quantified using quantile-quantile-plotting, as described by Reinhard^58^. This was implemented in a custom-made MATLAB script that additionally calculated the mean fluorescence background of the image, the mean fluorescence of the main population and of the subpopulation (of tc cells) (vs 2016a, MathWorks). To compare across ICE promoters, fluorescence values (F) were normalized by subtracting the image background (I) and divided by the same (i.e., (F–I)/I), and then averaged across three independent clones (data reported in Figure 2d). Significance of increased expression in the subpopulation of tc cells was tested on the three paired normalized values (one-sided paired t-test, H1 assuming that tc subpopulation expresses higher than main population).

### Time-lapse experiments

For time-lapse experiments, *P. knackmussii* B13 strains were precultured in LB, and then grown in MM with 3CBA and appropriate antibiotics, as described above. After 96 h incubation at 30°C, enough for cells to reach stationary phase and produce tc cells, the culture was diluted 100-fold in MM without carbon substrate added and transferred to microscope growth medium surfaces (“gel patches”). Four gel patches (volume each 0.13 ml, 1 mm thick and 6 mm ø) were cast in a microscope POC chamber, as described previously^58^. Gels contained 1% *w/v* agarose in MM with 0.1 mM 3CBA. Three patches were seeded with 6 μl of the 100-fold diluted cell suspension, left to dry at ambient air for 3-5 min in a laminar flow hood, then turned upside down and placed on round cover slip (42 mm ø, 0.17 mm thickness). A silicon spacer ring (1 mm thickness) was added and a second circular cover slip was put on top, after which the whole system was mounted in a rigid metal cast POC chamber and fixed with a metal ring (Reinhard, 2010). The POC chamber was incubated at 21 °C and images were taken (PhC, eGFP and mCherry) with a Plan Apo λ 100× 1.45 NA Oil objective during 48 h with intervals of 1 h at eight random positions using a Nikon Eclipse Ti-E Inverted Microscope, equipped with a Perfect Focus System (PFS) and pE-100 CoolLED illumination. In between imaging, the microscope lense was “parked” at the unseeded patch, in order to avoid illumination/heat damage to the cells. An in-house program written in Micro-Manager 1.4 was used to pilot the time-lapse experiments and record image series. Images were subsequently processed as described above, to automatically segment and position the cells across time-series, and to extract eGFP and mCherry per cell average fluorescence values. Cell identities given during segmentation were used to align corresponding eGFP and mCherry fluorescence profiles. Individual slopes were extracted as the linear regression of per cell average fluorescence increase (in arbitrary camera units h^-1^) during at least three consecutive time points as described previously^13^, which was then used to determine the onset of expression start relative to the incubation start (taken as t = 0 in Figure 5). Note that we did not take fluorescent protein maturation time into account for comparison of expression onsets between individual ICE promoters. Correlations of slopes were drawn using linear models.

### ICE*clc* transfer

ICE*clc* transfer assays were carried out, as described elsewhere^26^. In brief, *P. putida* ICE*clc* donors were cultured for 96 h in MM with 5 mM 3CBA (plus appropriate antibiotics to select for genetic constructions) to induce tc cell formation, whereas recipient cultures (*P. putida* UWCGC, gentamicin-resistant derivative of UWC1) were grown for 24 h in MM with 10 mM succinate and gentamicin. Recipient and donor cultures were mixed in a 1:2 volumetric ratio, respectively, in a total volume of 1 ml. Cells were harvested by centrifugation at room temperature for 1 min at 5000 × g, washed in 1 ml of MM without carbon substrate, centrifuged again and finally resuspended in 20 μl of MM. This mixture was deposited on top of a 0.2–μm cellulose acetate filter (Sartorius) placed on MM-agar containing 0.5 mM 3CBA, and incubated at 30°C for 48 h. Cells were then recovered from the filter by vortexing in 1 ml of MM, serially diluted in MM, and plated on selective plates. The same culture volumes of either donor or recipient alone were prepared and incubated similarly, to correct for the frequency of spontaneous background growth. Exconjugants were selected on MM agar plates with Gm and 3CBA (from transfer of ICE*clc*); donors on MM with 3CBA and Km, and recipients on MM with Gm and 10 mM succinate. Transfer frequencies are reported as the mean of the exconjugant colony forming units on MM-Gm-3CBA compared to that of the donor in the same assay (on MM-Km-3CBA).

### RNA-seq

Total sequencing of reverse-transcribed rRNA-depleted mRNAs (RNA-seq) was conducted on exponentially growing or ‘restimulated’ stationary phase cultures of *P. putida* UWC1 carrying wild-type ICE*clc* (strain 2737), *ICEclcΔmfsR* (strain 4322), or *ICEclcΔmfsRΔbisR* (strain 5553). Cultures were grown in fourfold replicates in MM with 5 mM 3CBA as described previously^59^ and harvested in exponential phase at a culture turbidity of 0.6 (at 600 nm). Four other replicate cultures were incubated for 96 h on 5 mM 3CBA (late stationary phase, to induce TCR), and then stimulated for 4.5 h by addition of 5 mM 3CBA (final concentration) to induce ICE excision and transfer. Cells were harvested by centrifugation as described, and total RNA was purified by hot phenol, DNAseI digestion, MiniElute cleanup, and depleted from rRNAs using the Ribo-Zero rRNA removal kit (EpiCentre)^59^. cDNA libraries were generated using a strand-specific ScriptSeq Complete Kit Bacteria protocol (Epicentre), indexed and sequenced on an Illumina HiSeq 2000 platform at the Lausanne Genomics Facilities with 101-nt single-end reads. Reads were cleaned and trimmed using TRIMMOMATIC^60^, then mapped, sorted and indexed using Bowtie2^61^ and Samtools^62^ under default settings, using the *P. putida* strain KT2440 chromosome (refseq NC_002947) and ICE*clc* (Genbank accession AJ617740.2) reconstructed genome sequence as reference. Mapped reads were counted with HTseq (version 0.11.2)^63^. Read counts were normalized using the PseudoReference Sample transformation^64^. In short, for each gene the geometric mean across all samples was calculated and used as the pseudoreference sample. Then for each sample, the read count of every gene was divided by its corresponding value in the pseudoreference. The median value of all those ratios of a given sample was used as the normalization factor for that sample. Data were log_2_(x+1) transformed in order to deal with zero values, before further clustering using *clustergram* as implemented in MATLAB (v. 2020a). For coverage plots, raw Htseq counts per position from a single replicate condition were read into MATLAB and plotted in a select window of 1000 bp covering the gene of interest. Plots were overlain for two conditions in Adobe Illustrator (v. 2020).

### Statistical methods

Mean background-normalized fluorescence expression values for different single-copy promoter fusions were compared between main and subpopulations of tc cells (*n* = 3 replicates with different clones, paired one-sided t-test). Slopes of fluorescence induction among single cells in time-lapse imaging were compared for paired promoters using linear regression models. Effects of BisDC induction on ICE promoter expression in absence of ICE*clc* in *P. putida* was tested on *n* = 3 independent replicates with different mini-Tn inserted promoter fusions, grown to stationary phase (48–96 h) on MM with 5 mM 3CBA. Because of strongly skewed fluorescence distributions we took here the 95^th^ percentile value to compare between strains carrying *bisDC* or with empty plasmid (paired one-sided t-test). Effects of InrR on the subpopulation sizes of cells expressing fluorescence from single-copy ICE promoters was tested on *n* = 3 independent replicates with different mini-Tn inserted promoter fusions, grown to stationary phase on MM with 5 mM 3CBA and sampled at 24, 48 and 72 h. Triplicate estimates of subpopulation sizes (by QQ-plotting) across the three strains were then compared in one-factorial ANOVA, implemented as the Bartlett test in *R*, followed by *aov* and Tukey multiple comparisons of means at 95% family-wise confidence level. The same procedure was followed to compare transfer rates among ICE-deletion variants.

## Supporting information

Sulser_Vucicevic_supplementary_information

## Data availability

The sequence read data belonging to the *P. putida* ICE transcriptomes are available from the Short Read Archives under project number **PRJNA784540**.

## Acknowledgements

This research has been supported by Swiss National Science Foundation grants 31003B_156926/1 and 31003A_175638 to JvdM. We thank Noémie Matthey for her help in initial parts of this project.

## Author contributions

SS, AV, VB, RM, FD, VS and NC performed experimental work. SS, AV, RM, NC and JvdM analyzed results. SS, NC and JvdM wrote the manuscript. JvdM was responsible for funding acquisition. All authors reviewed the manuscript and agreed to its submission.

