## Supplementary material for "A bistable orthogonal prokaryotic differentiation system underlying development of conjugative transfer competence": Sulser_Vucicevic_supplementary_information

Table S1: strain specifications
Table S2: used primer sets
Supplementary figure S1 - all PCR fragments
Supplementary figure S2 - RNAseq succinate
Supplementary figure S3 - Time lapse other constructs
Supplementary figure S4 – QQ-plots of BisDC induction
Supplementary figure S5 - Gene syntenies *orf88400-orf75419* region
Supplementary references

Supplementary table 1: Strain specifications.

| Strain number | Description | Remarks | Source or Reference |
|---|---|---|---|
| 78 | *P. knackmussii* B13 | Original host for ICE*clc* | [1] |
| 1291 | *P. putida* UWC1 | ICE*clc*-free control strain | [2] |
| 1343 | *P. knackmussii* B13 $P_{int}$-egfp, $Km^R$ | Single copy *intB13* promoter-egfp fusion insertion | [3] |
| 2058 | *P. knackmussii* B13 *inrR*$^{+/-}$ | Deletion of one copy of *inrR* | [4] |
| 2201 | *P. knackmussii* B13 *inrR*$^{-/-}$ | Deletion of both copies of *inrR* | [4] |
| 2397-98 | *P. knackmussii* B13 $P_{inR}$-egfp, $Km^R$ | Single copy *inrR* promoter-egfp fusion insertion | [4] |
| 2433 | *P. putida* UWC1 (ICE*clc-ΔinrR*) | By conjugation from 2201 | [4] |
| 2580-82 | *P. knackmussii* B13 $P_{int}$-egfp/$P_{inR}$-*mcherry*, $Km^R$ | Single copy bidirectional *intB13* promoter-egfp/*inrR* promoter-*mcherry* fusion insertion | [4] |
| 2737,2738 | *P. putida* UWC1 (ICE*clc*) | ICE*clc* element in 1291, $tRNA^{Gly}$ | [5] |
| 2744 | *P. putida* UWC1, mini-Tn7-$P_{tac}$-*mCherry*-$Gm^R$ | Constitutive mCherry expression, used as mating recipient | [5] |
| 2748 | *P. knackmussii* B13, mini-Tn5($P_{int}$-mcherry) $Tet^R$ | $P_{int}$ reporter in 78<br>262 bp fragment | [4] |
| 3044 | *E. coli* DH5α λpir | Host for propagation plasmids with R6K origin of replication | Victor de Lorenzo |
| 3734 | *E. coli* DH5α λpir, mini-Tn5 P(*orf81655*)-egfp, $Ap^R$ $Km^R$ | pBAM-egfp containing promoter region of *orf81655* | This study |
| 3737 | *E. coli* DH5α λpir, mini-Tn5 P(*orf81655*)-egfp $Km^R$ | pBAM-egfp containing promoter region of *orf81655* (fragment 4) | This study |
| 3741-3743 | *P. knackmussii* B13, mini-Tn5($P_{81655}$-egfp*)*, $Km^R$ | Single copy insertion of ($P_{81655}$-egfp*)* reporter of 3734 in 78 | This study |
| 3955 | *E. coli* DH5α λpir, mini-Tn5 P(*orf100952*)-egfp $Km^R$ | pBAM-egfp containing promoter region of *orf100952 (alpA)* | This study |
| 3956-58 | *P. knackmussii* B13, mini-Tn5 P(*orf100952*)-egfp $Km^R$ | $P_{alpA}$-egfp reporter from 3955 in 78 | This study |
| 3960 | *E. coli* DH5α λpir, mini-Tn5 P(*orf67231*)-egfp $Km^R$ | pBAM-egfp containing promoter region of *orf67231* | This study |
| 3961-63 | *P. knackmussii* B13, mini-Tn5 P(*orf67231*)-egfp $Km^R$ | $P_{orf67231-}$-egfp reporter from 3960 in 78 | This study |
| 3965 | *E. coli* DH5α λpir, mini-Tn5 P(*orf67800*)-egfp $Km^R$ | pBAM-egfp containing promoter region of *orf67800* | This study |
| 3966-68 | *P. knackmussii* B13, mini-Tn5 P(*orf67800*)-egfp $Km^R$ | $P_{orf67800}$-egfp reporter from strain 3965 in 78 | This study |
| 3970 | *E. coli* DH5α λpir, mini-Tn5 UR(*orf89746*)-egfp $Km^R$ | pBAM-egfp (strain 3726) containing upstream region of *orf89746* | This study |
| 3971-73 | *P. knackmussii* B13, mini-Tn5 UR(*orf89746*)-egfp $Km^R$ | $UR_{orf89746}$-egfp reporter from 3970 in strain 78 | This study |
| 3991 | *E. coli* DH5α λpir, mini-Tn5 UR(*orf66202*)-egfp $Km^R$ | pBAM-egfp containing upstream region of *orf66202* | This study |
| 3992-94 | *P. knackmussii* B13, mini-Tn5 UR(*orf66202*)-egfp $Km^R$ | $_{URorf66202}$-egfp reporter from 3991 in 78 | This study |
| 3996 | *E. coli* DH5α λpir, mini-Tn5 UR(*orf50240*)-egfp $Km^R$ | pBAM-egfp (strain 3726) containing short upstream region of *orf50240* | This study |
| 3997-99 | *P. knackmussii* B13, mini-Tn5 UR(*orf50240*)-egfp $Km^R$ | $_{URorf50240}$-egfp reporter from 3996 in 78 | This study |
| 4001 | *E. coli* DH5α λpir, mini-Tn5 P(*orf88400*)-egfp $Km^R$ | pBAM-egfp containing promoter region of *orf88400* | This study |

| | | | |
|---|---|---|---|
| **4002-04** | *P. knackmussii* B13, mini-Tn5 P(*orf88400*)-egfp $Km^R$ | $P_{orf88400}$-egfp reporter from 4001 in 78 | This study |
| **4006** | *E. coli* DH5α λpir, mini-Tn5 P(*orf101284*)-egfp $Km^R$ | pBAM-egfp containing promoter region of *orf101284* (*bisR*) | [6] |
| **4007-09** | *P. knackmussii* B13, mini-Tn5 P(*orf101284*)-egfp $Km^R$ | $P_{bisR}$-egfp reporter from 4006 in 78 | This study |
| **4011** | *E. coli* DH5α λpir, mini-Tn5 UR(*orf100033*)-egfp $Km^R$ | pBAM-egfp containing (short) upstream region of *orf100033* | This study |
| **4012-14** | *P. knackmussii* B13, mini-Tn5 UR(*orf100033*)-egfp $Km^R$ | $UR_{100033}$-egfp reporter from 4011 in 78 | This study |
| **4064** | *E. coli* DH5α λpir, mini-Tn5 UR(*orf84835*)-egfp $Km^R$ | pBAM-egfp containing upstream region of *orf84835* | This study |
| **4065-67** | *P. knackmussii* B13, mini-Tn5 UR(*orf84835*)-egfp $Km^R$ | $_{URorf84835}$-egfp reporter of 4064 in 78 | This study |
| **4096-98** | *P. knackmussii* B13, mini-Tn5 P(*orf81655_4*)-egfp , $P_{int}$-*eChe*, Tc,$Km^R$ | $P_{orf81655}$-egfp fragment–4 reporter of 3737 in 2748 | This study |
| **4122** | *E. coli* DH5α λpir, mini-Tn5 UR(*orf62755*)-egfp $Km^R$ | pBAM-egfp containing upstream region of *orf62755* | This study |
| **4123-25** | *P. knackmussii* B13, mini-Tn5 UR(*orf62755*)-egfp $Km^R$ | $UR_{62755}$-egfp reporter of 4122 in 78 | This study |
| **4126-28** | *P. knackmussii* B13, mini-Tn5 P(*orf100952*)-egfp $Km^R$, mini-Tn5($P_{int}$-mcherry) $Tet^R$ | $P_{orf100952}$-egfp reporter in 2748 (double reporter) | This study |
| **4129-31** | *P. knackmussii* B13, mini-Tn5 P(*orf67231*)-egfp $Km^R$, mini-Tn5($P_{int}$-mcherry) $Tet^R$ | $P_{orf67231}$-egfp reporter in 2748 (double reporter) | This study |
| **4322** | *P. putida* UWC1 ICE*clc* Δ*mfsR* | Deletion of *mfsR*, used for RNAseq | [7] |
| **4323-25** | *P. knackmussii* B13, mini-Tn5 UR(*orf100033*)-egfp $Km^R$, mini-Tn5($P_{int}$-mcherry) $Tet^R$ | $UR_{orf100033}$-egfp reporter in 2748 | This study |
| **4326-28** | *P. knackmussii* B13, mini-Tn5 P(*orf101284*)-egfp $Km^R$, mini-Tn5($P_{int}$-mcherry) $Tet^R$ | $P_{orf101284}$-egfp reporter in 2748 (double reporter) | This study |
| **4369-71** | *P. knackmussii* B13, mini-Tn5 P(*orf88400*)-egfp $Km^R$, mini-Tn5($P_{int}$-mcherry) $Tet^R$ | $P_{orf88400}$-egfp reporter in 2748 (double reporter) | This study |
| **4764-66** | *P. knackmussii* B13 *inrR*$^{+/-}$, mini-Tn5 P(*orf88400*)-egfp $Km^R$ | $P_{orf88400}$-egfp reporter in strain 2058 | This study |
| **4767-69** | *P. knackmussii* B13 *inrR*$^{-/-}$, mini-Tn5 P(*orf88400*)-egfp $Km^R$ | $P_{orf88400}$-egfp reporter in strain 2201 | This study |
| **4770-72** | *P. knackmussii* B13 *inrR*$^{+/-}$, mini-Tn5 P(*orf100952*)-egfp $Km^R$ | $P_{orf100952}$-egfp reporter in strain 2058 | This study |
| **4773-75** | *P. knackmussii* B13 *inrR*$^{-/-}$, mini-Tn5 P(*orf100952*)-egfp $Km^R$ | $P_{orf100952}$-egfp reporter in strain 2201 | This study |
| **4776-78** | *P. knackmussii* B13 *inrR*$^{+/-}$, mini-Tn5 P(*orf101284*)-egfp $Km^R$ | $P_{orf101284}$-egfp reporter in strain 2058 | This study |
| **4779-81** | *P. knackmussii* B13 *inrR*$^{-/-}$, mini-Tn5 P(*orf101284*)-egfp $Km^R$ | $P_{orf101284}$-egfp reporter in strain 2201 | This study |
| **4782-84** | *P. knackmussii* B13 *inrR*$^{+/-}$, mini-Tn5 P(*orf81655*)-egfp $Km^R$ | $P_{orf81655}$-egfp reporter in strain 2058 | This study |
| **4785-87** | *P. knackmussii* B13 *inrR*$^{-/-}$, mini-Tn5 P(*orf81655*)-egfp $Km^R$ | $P_{orf81655}$-egfp reporter in strain 2201 | This study |
| **4822-24** | *P. knackmussii* B13 *inrR*$^{+/-}$, mini-Tn5 P(*orf67231*)-egfp $Km^R$ | $P_{orf67231}$-egfp reporter in strain 2058 | This study |
| **4826-28** | *P. knackmussii* B13 *inr*$^{-/-}$, mini-Tn5 P(*orf67231*)-egfp $Km^R$ | $P_{orf67231}$-egfp reporter in strain 2201 | This study |
| **4852-54** | *P. putida* UWC1 *ΔmfsR* P(*orf88400*)-egfp $Km^R$ | $P_{orf88400}$-egfp reporter in 4322 | This study |
| **4855-57** | *P. putida* UWC1 *ΔmfsR* P(*orf100952*)-egfp $Km^R$ | $P_{orf100952}$-egfp reporter in 4322 | This study |
| **4858-60** | *P. putida* UWC1 *ΔmfsR* P(*orf101284*)-egfp $Km^R$ | $P_{orf101284}$-egfp reporter in 4322 | This study |
| **4882-84** | *P. putida* UWC1 *ΔmfsR* P(*orf81655*)-egfp $Km^R$ | $P_{orf81655}$-egfp reporter in 4322 | This study |

| | | | |
|---|---|---|---|
| **5284** | *E. coli* DH5a-lambda pir P(orf58432)-egfp, $Ap^R$ | $P_{orf58432}$-egfp reporter in pBAM | This study |
| **5294-96** | *P. putida* UWC1 *ΔmfsR* P(*orf67231*)-egfp $Km^R$ | $P_{orf67231}$-egfp reporter in 4322 | This study |
| **5337-39** | *P. knackmussii* B13 mini-Tn5 P(*orf58432*)-egfp $Km^R$ | P(*orf58432*)-egfp reporter from 5284 in 78 | This study |
| **5501** | *P. putida* UWC1 miniTn7::P(*int*)-egfp $Gm^R$, pME6032 $Tet^R$ | $P_{int}$-egfp, empty plasmid pME6032 | [6] |
| **5502** | *P. putida* UWC1 miniTn7::P(*inR*)-egfp $Gm^R$, pME6032 $Tet^R$ | $P_{inR}$-egfp, empty plasmid pME6032 | [6] |
| **5503** | *P. putida* UWC1 miniTn7::P(*alpA*)-egfp $Gm^R$, pME6032 $Tet^R$ | $P_{alpA}$-egfp, empty plasmid pME6032 | [6] |
| **5553** | *P. putida* UWC1 clc5, ICE*clc* Δ*mfsR* Δ*bisR* | Derivative of 4322 with additional *bisR* deletion, used for RNAseq | This study |
| **5719, 5725, 5731** | *P. putida* UWC1 $P_{int}$-*mcherry*/$P_{inR}$-*egfp*, pME6032, $Km^R$, $Tc^R$ | Control for direction induction of Pint,PinR by BisDC | This study |
| **5929** | *E. coli* DH5α λpir, mini-Tn5 UR(*orf97571*)-egfp $Km^R$ | pBAM-egfp containing upstream region of *orf97571* | This study |
| **5930-32** | *P. knackmussii* B13 mini-Tn5-UR(*orf97571*)-egfp insertion | UR(*orf97571)-egfp* reporter of 5929 in strain 78 | This study |
| **5933** | *E. coli* DH5α λpir, mini-Tn5 UR(*orf100033*)-egfp $Km^R$ | pBAM-egfp containing upstream region of *orf100033* | This study |
| **5934-36** | *P. knackmussii* B13 mini-Tn5-UR(*orf100033*)-egfp insertion | UR(*orf100033)-egfp* reporter of 5933 in strain 78 | This study |
| **5937-39** | *P. knackmussii* B13 *inrR*$^{-/-}$, mini-Tn5 P(inR)-egfp $Km^R$ | PinR-egfp reporter from 2012 into 2201 | This study |
| **5940-42** | *P. knackmussii* B13 miniTn5-$P_{int}$-*mcherry*; mini-Tn5-P(*orf58432*)-egfp insertion | P(*orf58432)-egfp* reporter of 5248 in strain 2748 | This study |
| **6059-61** | *P. putida* UWC1 $P_{int}$-*mcherry*/$P_{inR}$-*egfp*, pMEbisDC, $Km^R$, $Tc^R$ | Strain 5690 with plasmid from 6055 | This study |
| **6065** | *P. putida* UWC1 miniTn7::P(*alpA*)-egfp $Gm^R$, pMEbisDC $Tet^R$ | $P_{alpA}$-egfp, bisDC expression from pMEbisDC | [6] |
| **6077** | *P. putida* UWC1 miniTn7::P(*int*)-egfp $Gm^R$, pMEbisDC $Tet^R$ | $P_{int}$-egfp, bisDC expression from pMEbisDC | [6] |
| **6178-80** | *P. knackmussii* B13 *inrR*$^{+/-}$, mini-Tn5 P(inR)-egfp $Km^R$ | PinR-egfp reporter from 2012 into 2058 | This study |
| **6298-6300** | *P. putida* UWC1 miniTn5::P(*orf67231*)-egfp $Km^R$, pME6032 $Tet^R$ | $P_{orf67231}$-egfp reporter in 1291, empty plasmid pME6032 | This study |
| **6301-03** | *P. putida* UWC1 miniTn5::P(*orf67231*)-egfp $Km^R$, pMEbisDC $Tet^R$ | $P_{orf67231}$-egfp reporter in 1291, bisDC expression from pMEbisDC | This study |
| **6304-06** | *P. putida* UWC1 miniTn5::UR(*orf89746*)-egfp $Km^R$, pME6032 $Tet^R$ | $UR_{orf89746}$-egfp reporter in 1291, empty plasmid pME6032 | This study |
| **6307-09** | *P. putida* UWC1 miniTn5::UR(*orf89746*)-egfp $Km^R$, pMEbisDC $Tet^R$ | $UR_{orf89746}$-egfp reporter in 1291, bisDC expression from pMEbisDC | This study |
| **6310-12** | *P. putida* UWC1 miniTn5::P(*orf88400*)-egfp $Km^R$, pME6032 $Tet^R$ | $P_{orf88400}$-egfp reporter in 1291, empty plasmid pME6032 | This study |
| **6313-15** | *P. putida* UWC1 miniTn5::P(*orf88400*)-egfp $Km^R$, pMEbisDC $Tet^R$ | $P_{orf88400}$-egfp reporter in 1291, bisDC expression from pMEbisDC | This study |
| **6856** | *P. putida* UWC1 miniTn7::P(*inR*)-egfp $Gm^R$, pMEbisDC $Tet^R$ | $P_{inR}$-egfp, bisDC expression from pMEbisDC | [6] |
| **7150** | *E. coli* DH5α λpir, *mini-Tn5 (P50240)-egfp* $Km^R$ | pBAM-egfp containing long promoter region of *orf50240* | This study |
| **7177-79** | *P. putida* UWC1 ICEclc5 miniTn5::P(*orf50240*)-egfp $Km^R$ | $P_{orf50240}$-egfp reporter of 7150 in 2737 | This study |

| | | | |
|---|---|---|---|
| **7181-82** | *P. putida* UWC1 miniTn5::P(*orf50240*)-egfp $Km^R$, pMEbisDC $Tet^R$ | $P_{orf50240}$-egfp reporter (short version) of 7149 in 1291, bisDC expression from pMEbisDC | This study |
| **7183-84** | *P. putida* UWC1 miniTn5::P(*orf50240*)-egfp $Km^R$, pMEbisDC $Tet^R$ | $P_{orf50240}$-egfp reporter of 7150 in 1291, bisDC expression from pMEbisDC | This study |
| **7205-06** | *P. putida* UWC1 miniTn5::P(*orf50240*)-egfp, $Km^R$, pME6032 $Tet^R$ | $P_{orf50240}$-egfp reporter from 7150 in 1291, empty plasmid pME6032 | This study |
| **7222-24** | *P. knackmussii* B13 *inrR*$^{+/-}$, mini-Tn5 P(*orf50240*)-egfp $Km^R$ | $P_{orf50240}$ -egfp reporter from 7150 in 2058 | This study |
| **7227-29** | *P. knackmussii* B13 *inrR*$^{-/-}$, mini-Tn5 P(*orf50240*)-egfp $Km^R$ | $P_{orf50240}$ -egfp reporter from 7150 in 2201 | This study |
| **7310** | *E. coli* DH5$\alpha$ $\lambda$pir, *mini-Tn5 (UR84835)-egfp* $Km^R$ | pBAM-egfp (strain 3726) containing upstream region of *orf84835* | This study |
| **7311** | *E. coli* DH5$\alpha$ $\lambda$pir, *mini-Tn5 (UR89247)-egfp* $Km^R$ | pBAM-egfp (strain 3726) containing upstream region of *orf89247* | This study |
| **7330-32** | *P. putida* UWC1 ICEclc5 miniTn5::UR(*orf84835*)-egfp $Km^R$ | $_{URorf84835}$-egfp reporter from 7310 in 2737 | This study |
| **7334-36** | *P. putida* UWC1 ICEclc5 miniTn5::UR(*orf89247*)-egfp $Km^R$ | $_{URorf89247}$-egfp reporter from 7311 in 2737 | This study |
| **7378-80** | *P. putida* UWC1 ICEclc6 Δ81655-75419 mini-Tn5(PinR-egfp/Pint-mcherry) $Km^R$ | $P_{inR}$-egfp/ $P_{int}$-mcherry reporter in 2738 carrying ICE*clc* deleted for orfs *81655-75419* | This study |
| **7381-83** | *P. putida* UWC1 ICEclc6 Δ88400-84388 mini-Tn5(PinR-egfp/Pint-mcherry) $Km^R$ | $P_{inR}$-egfp/ $P_{int}$-mcherry reporter in 2738 carrying ICE*clc* deleted for orfs *88400-84388* | This study |
| **7384-86** | *P. putida* UWC1 ICEclc5 mini-Tn5(PinR-egfp/Pint-mcherry) $Km^R$ | $P_{inR}$-egfp/ $P_{int}$-mcherry reporter in 2737 | This study |
| **7387-89** | *P. putida* UWC1 ICEclc6 Δ74436-68241 mini-Tn5(PinR-egfp/Pint-mcherry) $Km^R$ | $P_{inR}$-egfp/ $P_{int}$-mcherry reporter in 2738 carrying ICE*clc* deleted for orfs *74436-68241* | This study |
| **7418** | *E. coli* DH5$\alpha$ $\lambda$pir, *mini-Tn5::UR(73676)-egfp* $Km^R$ | pBAM-egfp containing upstream region of *orf73676* | This study |
| **7423-25** | *P. putida* UWC1 ICEclc5 miniTn5::UR(*orf73676*)-egfp $Km^R$ | $UR_{orf73676}$-egfp reporter from 7418 in 2737 | This study |
| **7476-78** | *P. putida* UWC1 miniTn5::P(*orf81655 fgt4*)-egfp $Km^R$, pME6032 $Tet^R$ | $P_{orf81655}$-egfp reporter from 3448 in 1291, empty plasmid pME6032 | This study |
| **7479-81** | *P. putida* UWC1 miniTn5::P(*orf81655 fgt4*)-egfp $Km^R$ , pMEbisDC $Tet^R$ | $P_{orf81655}$-egfp reporter from 3448 in 1291, bisDC expression from pMEbisDC | This study |
| **7558-60** | *P. putida* UWC1 miniTn5::P(*orf50240*)-egfp, miniTn5::$P_{int}$-*mcherry* $Km^R$ , $Tet^R$ | Double reporter fusion, miniTn5 of strain 2707 in 7178 | This study |

Supplementary table S2: Primers used.

| Number | Sequence 5'-3' | Target | Restriction site | Objective |
|---|---|---|---|---|
| 051005 | CAAGAAGGACCATGTGGTC | eGFP gene on pBAM | - | Construct verification |
| 060605 | TTTTTTGAATTCGCGCAATCACCGATCGCGCAT | *inr*R gene on ICE*clc* (94688-94708) | - | Amplification of inrR gene |
| 060606 | TTTTTTTCTAGAATGAGCGATCTGAACCAACCG | *inr*R gene on ICE*clc* (95201-95220) | - | |
| 070418 | CAGGAAACAGCTATGACC | Universal M13 primer | - | Construct verification |
| 090803 | TTTTGAATTCCTTGCCAAGGTCGGGGTC | $P_{int}$ (32-49) | *EcoRI* | *Amplification of Pint-mcherry reporter system* |
| 110404 | CTTCAGCGTCATAATGGC | mcherry gene | - | |
| 111201 | TTTTTGGATCCTTCGCTGGAACAGAGAGAGCAT | *orf50240* of ICE*clc* (52071-52093) | *BamH*I | *Short upstream region of 50240* |
| 111202 | TTTTTTCTAGAGATGCTCTCCAGAGTCCAAGAAT | *orf50240* of ICE*clc* (52307-52329) | *Xba*I | |
| 111203 | TTTTTTCTAGATTCGCTGGAACAGAGAGAGCAT | *orf52324-53196* of ICE*clc* (52071-52093) | *Xba*I | *Opposite orientation* |
| 111204 | TTTTTGGATTCGATGCTCTCCAGAGTCCAAGAAT | *orf52324-53196* of ICE*clc* (52307-52329) | *BamH*I | |
| 111205 | TTTTTGGATCCAAAGACATGGCGAACCTCCGGA | *orf53587-58432* of ICE*clc* (58925-58946) | *BamH*I | *Upstream region of 58432* |
| 150803 | TTTTTTCTAGACCTGCTTGATCGCCA | *orf53587-58432* of ICE*clc* (59252..59266) | *Xba*I | |
| 111207 | TTTTTGGATCCCCAGATGTCCATCATTGTGCT | *orf59110-62755* of ICE*clc* (63082-63102) | *BamH*I | *Upstream region of 62755* |
| 111208 | TTTTTTCTAGAATTTCCAGCAACGAAGCCGTA | *orf59110-62755* of ICE*clc* (63383-63403) | *Xba*I | |
| 111209 | TTTTTGGATCCGATCGTGCGGAACACCCA | *orf63176-66202* of ICE*clc* (66439-66456) | *BamH*I | *Upstream region of 66202* |
| 111210 | TTTTTTCTAGAATTGCTGGTTGTGGCGTCGAT | *orf63176-66202* of ICE*clc* (66785-66805) | *Xba*I | |
| 111211 | TTTTTGGATCCATCACGAGAAAAGTGGGCAC | *orf66625-67231* of ICE*clc* (67562-67581) | *BamH*I | *Upstream region of 67231* |
| 111212 | TTTTTTCTAGAACATAGACCACTCAACGAGA | *orf66625-67231* of ICE*clc* (67901-67920) | *Xba*I | |
| 111213 | TTTTTTCTAGAATCACGAGAAAAGTGGGCAC | *orf67800* of ICE*clc* (67562-67581) | *Xba*I | *Upstream region of 67800* |
| 111214 | TTTTTGGATCCACATAGACCACTCAACGAGA | *orf67800* of ICE*clc* (67901-67920) | *BamH*I | |
| 120203 | TTTTTGGATCCGGACGGGCTCCTTGGAAAAG | *orf85934-88400* of ICE*clc* (88619-88638) | *BamH*I | *Upstream region of 88400* |
| 120204 | TTTTTTCTAGAGACCCTCATTACATCGACATGAC | *orf85934-88400* of ICE*clc* (89229-89252) | *Xba*I | |
| 120205 | TTTTTGGATCCATGTCGGGTCTCCTGTTCGT | *orf89247-89746* of ICE*clc* (91351-91370) | *BamH*I | *Upstream region of 89746* |
| 120206 | TTTTTTCTAGATTCCTGACCCGACCCCGTTC | *orf89247-89746* of ICE*clc* (91876-91895) | *Xba*I | |
| 120207 | TTTTTGGATCCTAATAGGACGACAACGTGGG | *orf96323-100033* of ICE*clc* (100771-100790) | *BamH*I | *Upstream region of 100033* |
| 120208 | TTTTTTCTAGATGATGCGTGCCGGCAAGTTC | *orf96323-100033* of ICE*clc* (101050-101069) | *Xba*I | |
| 120209 | TTTTTGGATCCTTCATCGAGACGCAAGATGC | *orf100952* of ICE*clc* (101113-101132) | *BamH*I | *Upstream region of 100952* |
| 120210 | TTTTTTCTAGAATTACCGATCGCACGCTGCAA | *orf100952* of ICE*clc* (101337-101357) | *Xba*I | |
| 120211 | TTTTTGGATCCATGCTCCGTCTCCTTCCAGGA | *orf101284* of ICE*clc* (102043-102063) | *BamH*I | *Upstream region of 101284* |
| 120212 | TTTTTTCTAGACGCAGTCGTCACAACGTCAT | *orf101284* of ICE*clc* (102655-102674) | *Xba*I | |
| 130701 | TTTTAAGCTTCGAGGTGTGAAGGTCGAAG | *81655up_for (82531-82549)* | *HindIII* | |

| | | | | |
|---|---|---|---|---|
| 130702 | TTTTTTGGATCCTGCGCATGGACAGGCCATT | *81655up_rev (83337-83355)* | *BamHI* | *Upstream region for deletion of 81655-75419* |
| 130703 | TTTTTTTCTAGATCAATCCCGAGCCAGCTTC | *75419down_for (74442-74460)* | *XbaI* | *Downstream region for deletion of 81655-75419* |
| 130704 | TTTTTAAGCTTCGCCGCCGGTTTCTTGTTA | *75419down_rev (75383-75401)* | *HindIII* | |
| 130705 | TTTTAAGCTTGAAACTCTCCTTGGGATAG | *74436up_for (75312-75330)* | *HindIII* | *Upstream region for deletion of 74436-68241* |
| 130706 | TTTTTTGGATCCTCAAGGACATCGGTGGCAA | 74436up_rev (76230-76248) | BamHI | |
| 130707 | TTTTTTTCTAGACAGCCGCGGGTAATCGAAG | *68241down_for (67390-67408)* | *XbaI* | *Downstream region for deletion of 74436-68241* |
| 130708 | TTTTTTAAGCTTACCGGCAGCGCAGGAAGTA | 68241down_rev (68219-68237) | HindIII | |
| 210611 | AGGGATAACAGGGTAATCTGAATTCCCCACCAGTTCGCCTCGGCAT | *D88400-84388UP.f* (89622-89642) | EcoRI | *Upstream region for deletion of 88400-84388* |
| 210612 | TCAGGCTTGCTGGGACATGGACGGGCT | *D88400-84388UP.r* (88604-88627) | - | |
| 210613 | AGCCCGTCCATGTCCCAGCAAGCCTGAAGCAGCAGCACGTCCTGCTCA | *D88400-84388DW.f* (84323-84343) | - | *Downstream region for deletion of 88400-84388* |
| 210614 | GAAGCTTGCATGCCTGCAGGTCGACGGAGTTGCAGCCGTGGCGCGAT | *D88400-84388DW.r* (83314-8335) | SalI | |
| 200901 | AATCAGAATTCGAGCTCGCCCTAGTGGCGGGCCGACCGGGA | *P50240.F* (52104-52123) | - | *Long upstream region of 50240* |
| 200903 | TTCGAGGCATGCCTGCAGCCCCGATCACACAGTCGTGGAA | *P50240.R2* (52743-52761) | - | |
| 210615 | AATCAGAATTCGAGCTCGCCCGAGGAAGGACAAGGGATTCC | *P84835.f* (85666-85685) | - | *Upstream region of 84835* |
| 210616 | TTCGAGGCATGCCTGCAGCCCGCGCCACGCACCATTCTTCCT | *P84835.R2* (86290-86310) | - | |
| 210617 | AATCAGAATTCGAGCTCGCCCGAAATCGACCGGGTGGCGACAT | *P89247.f* (89517-89538) | - | *Upstream region of 89247* |
| 210618 | TTCGAGGCATGCCTGCAGCCCCAAGGCAATCGGCAATTCCAT | *P89247.r* (89830-89850) | - | |
| 210619 | AATCAGAATTCGAGCTCGCCCCTCTCGGGCCACGAGCGGCA | *P73676.f* (74320-74339) | - | *Upstream region of 73676* |
| 210620 | TTCGAGGCATGCCTGCAGCCCGTGGACGCCTGGTCGATCGAT | *P73676.r* (75123-75143) | - | |

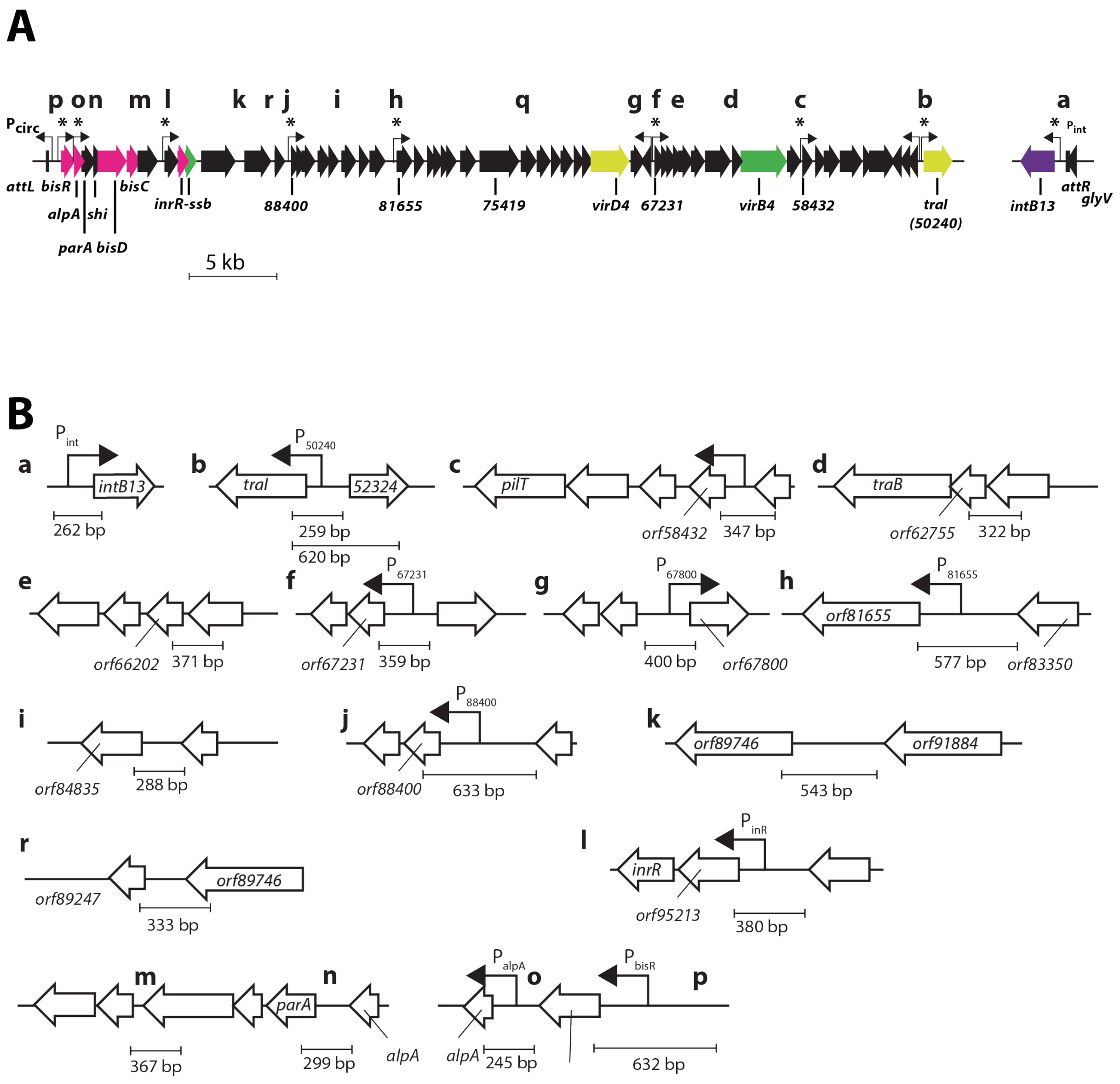


Supplementary figure 1. Cloned and tested upstream/promoter fragments of ICE core genes. **A** General overview of the ICE core region. **B** Individual selected promoter/upstream fragments and their sizes (small cap letters corresponding to fragment indications in the main text).

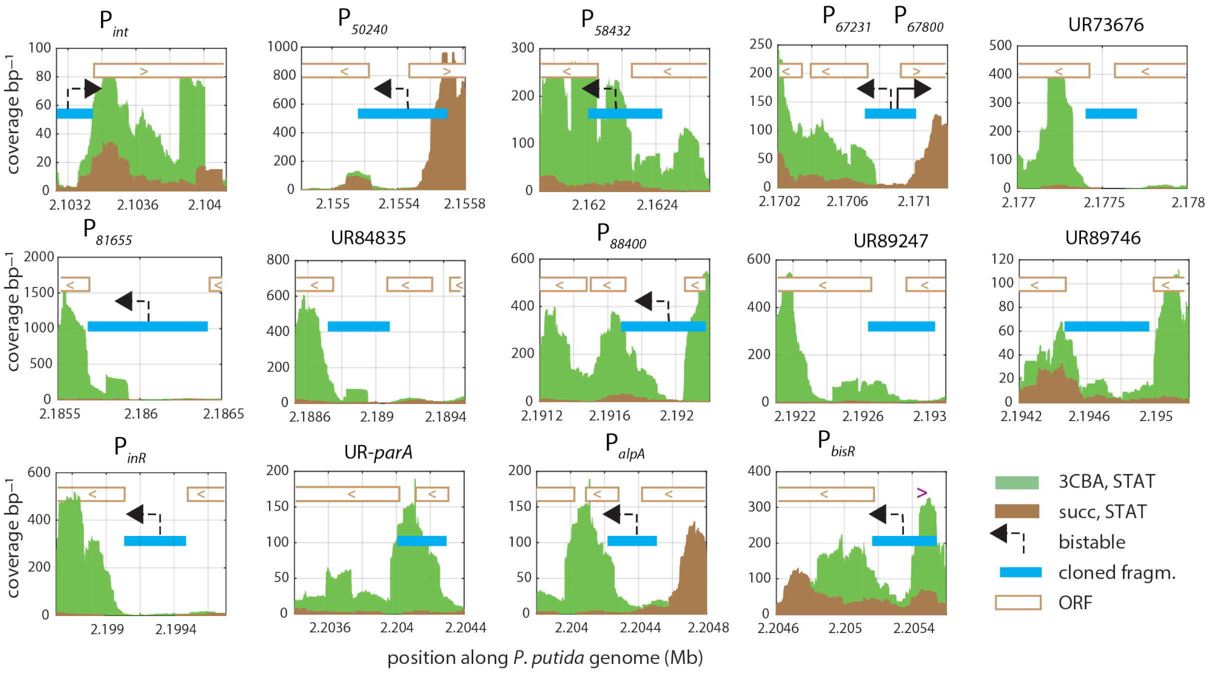


Supplementary figure 2. Read coverage of ICE*clc* transcripts in *P. putida* ICE*clc* in stationary phase conditions after growth with 3CBA (green) or succinate (brown) as carbon substrate. Plots show read coverage per basepair position from RNA-seq (for a single representative replicate) of the indicated conditions, plotted for the relevant ICE*clc* region on the x-axis (in Mbp). Blue bars with black arrows point to cloned fragments for single cell expression studies and their assumed promoter content (dotted arrow representing the bistably expressed identified transfer competence promoters on the ICE). Open directional bars correspond to relevant coding regions on ICE*clc*.

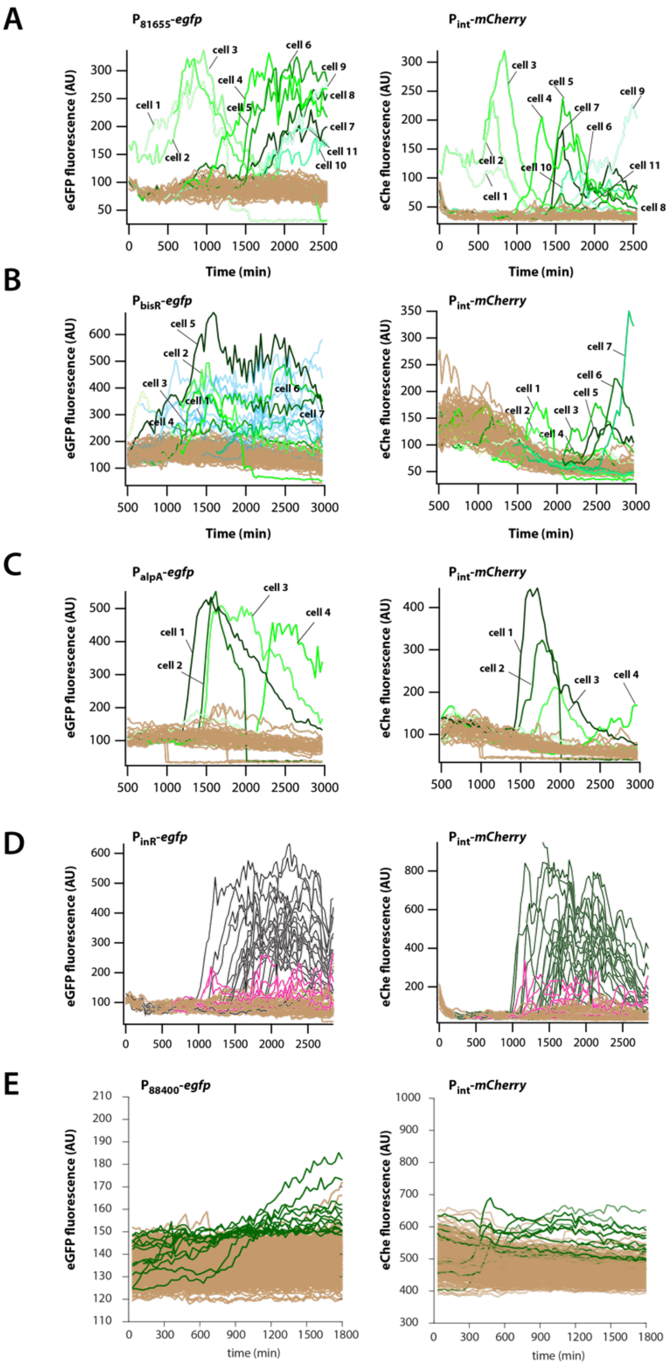


Supplementary figure 3. Time-lapse fluorescence of individual cells of surface-grown *P. knackmussii* B13 with indicated single-copy inserted promoter-fluorescent reporter constructs (**A-E**, one replicate strain for each); each line corresponding to an individual cell traced over time. Line traces are colored according to attribution of an individual cell as transfer competent (colored) or not (brown), based on quantile-quantile plotting at time point 20 h in both channels (plus retracing its previous history). P$_{int}$-*mcherry* insertions in same position on the genome for all constructs.

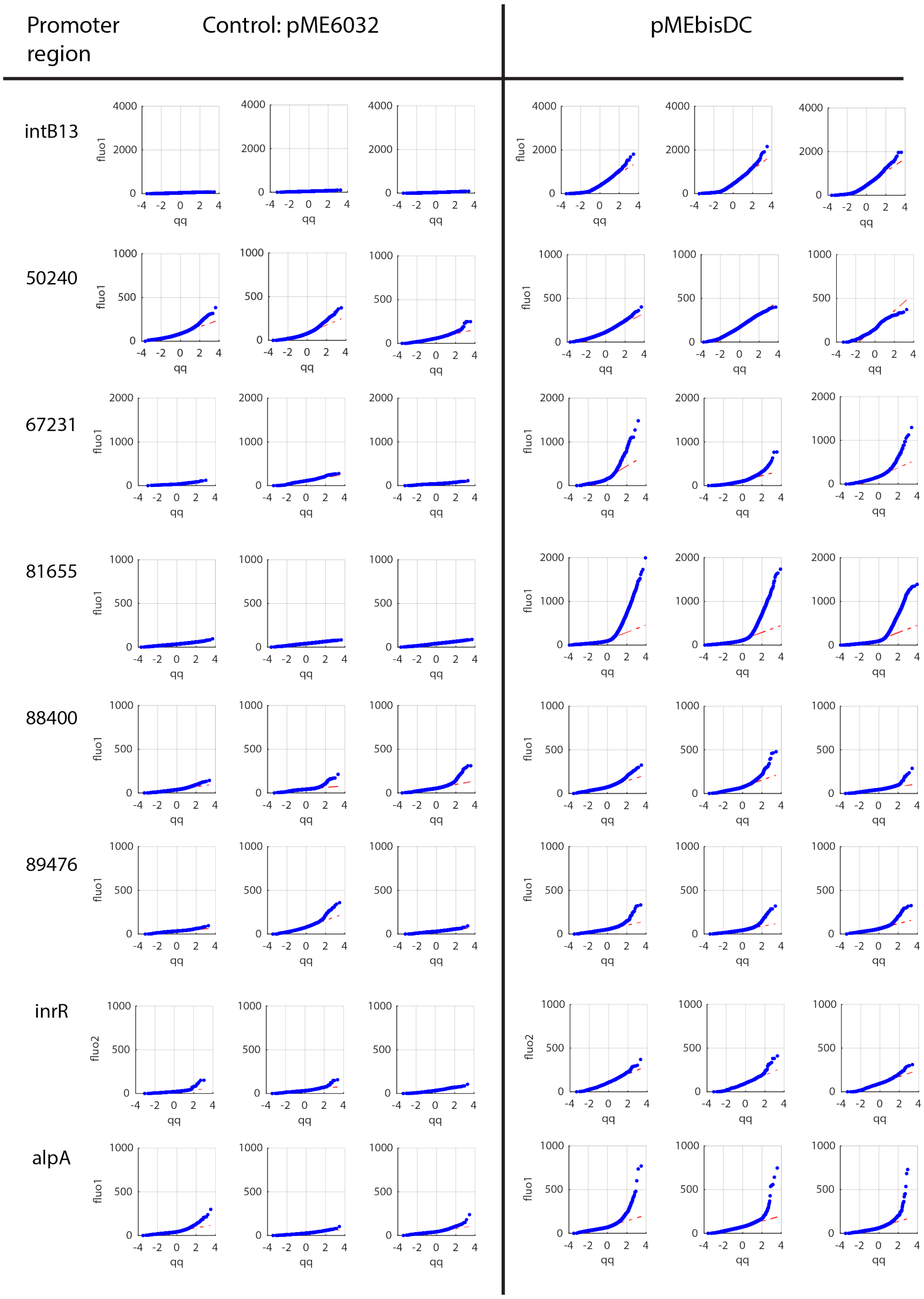


Supplementary figure 4. Cell fluorescence distribution from indicated single copy promoter-reporter fusions in *P. putida* without ICE*clc*, but induced or not for production of the BisDC activator complex (pMEbisDC). Cells sampled in stationary phase after growth on succinate. Comparisons are the same *P. putida* reporter strains but with empty plasmid (pME6032). Cell fluorescence distributions are plotted as their expected versus observed quantile; each plot showing a single biological replicate with independent reporter gene insertion position, grouped from n=10 images per sample. Each dot corresponds to a single segmented cell observation. Note the strongly tailed distributions for some constructs.

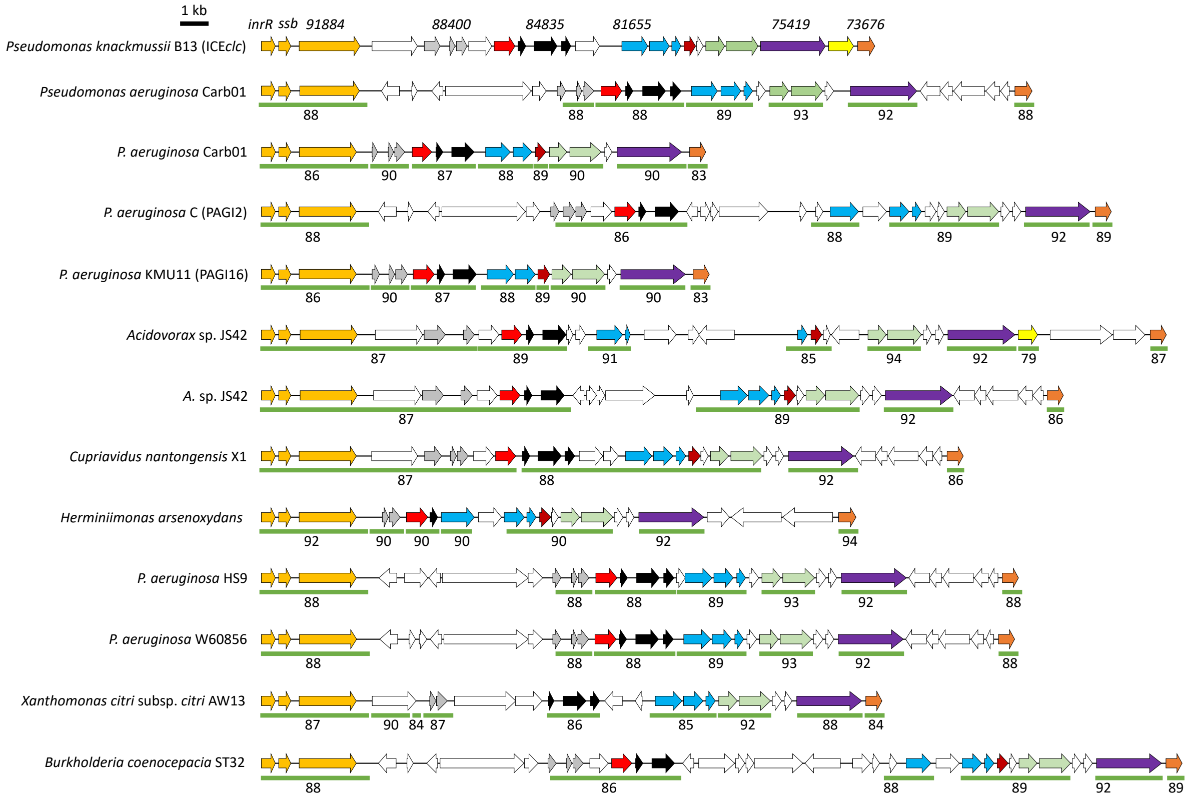


Supplementary figure 5. ICE*clc* gene and gene synteny conservation to putative ICE in genomes of other Gamma- and Betaproteobacteria. Regions are aligned to the gene cluster containing *inrR* and *ssb* (ochre), and then emphasize the conserved regions with unknown functions *orf88400* – *orf81655*. Ortholog genes are colored similarly. Arrows indicate the corresponding open reading frame length and orientation. Numbers below represent the percent nucleotide identity to ICE*clc*. White open arrows point to open reading frames not generally conserved with ICE*clc*.
